# Enhancing the spectral range of plant and bacterial Light-Harvesting pigment-protein complexes with various synthetic chromophores incorporated into lipid vesicles

**DOI:** 10.1101/2022.08.02.502557

**Authors:** Ashley M. Hancock, David J. K. Swainsbury, Sophie A. Meredith, Kenichi Morigaki, C. Neil Hunter, Peter G. Adams

## Abstract

The Light-Harvesting (LH) pigment-protein complexes found in photosynthetic organisms have the role of absorbing solar energy with high efficiency and transferring it to reaction centre complexes. LH complexes contain a suite of pigments that each absorb light at specific wavelengths, however, the natural combinations of pigments within any one protein complex do not cover the full range of solar radiation. Here, we provide an in-depth comparison of the relative effectiveness of five different organic “dye” molecules (Texas Red, ATTO, Cy7, DiI, DiR) for enhancing the absorption range of two different LH membrane protein complexes (the major LHCII from plants and LH2 from purple phototrophic bacteria). Proteoliposomes were self-assembled from defined mixtures of lipids, proteins and dye molecules and their optical properties were quantified by absorption and fluorescence spectroscopy. Both lipid-linked dyes and alternative lipophilic dyes were found to be effective excitation energy donors to LH protein complexes, without the need for direct chemical or generic modification of the proteins. The Förster theory parameters (e.g., spectral overlap) were compared between each donor-acceptor combination and found to be good predictors of an effective dye-protein combination. At the highest dye-to-protein ratios tested (over 20:1), the effective absorption strength integrated over the full spectral range was increased to ~180% of its natural level for both LH complexes. Lipophilic dyes could be inserted into pre-formed membranes although their effectiveness was found to depend upon favourable physicochemical interactions. Finally, we demonstrated that these dyes can also be effective at increasing the spectral range of surface-supported models of photosynthetic membranes, using fluorescence microscopy. The results of this work provide insight into the utility of self-assembled lipid membranes and the great flexibility of LH complexes for interacting with different dyes.

## Introduction

In most photosynthetic organisms, light-harvesting (LH) protein complexes have the primary function of absorbing solar energy with extremely high efficiency and transferring it to reaction centre (RC) complexes where photochemistry transduces this energy to a chemical form [1, 2]. LH complexes in plant and bacterial membranes act as “peripheral antennas” for the RC and contain high concentrations of carotenoids and chlorophyll(Chl)-type pigments for light absorption [3, 4]. The biochemical process of producing Chl molecules is complex and energy consuming [5], which limits the number of chemical variants of (bacterio)chlorophyll (B)Chl molecules that organisms can produce. Phototrophs appear to have evolved to produce appropriate LH complexes for their own ecological niche and do not attempt to cover the entire solar spectrum [6]. Instead, a particular LH complex will contain multiple copies of a few specific pigment types, with these arranged via precise protein-pigment interactions, resulting in a characteristic absorption spectrum. Such spectra reveal the wavelengths of light where absorption is highly effective but also highlight the “spectral gaps” where there is minimal light absorption for any given LH complex [7, 8]. Strong dipole-dipole and energetic coupling between closely-spaced (B)Chl molecules can allow excitation energy to be delocalised across multiple pigments and promote highly efficient transfer of excitation energy within the network of LH and RC protein complexes [9]. These transfers may be modelled by a combination of modified Redfield theory for strongly coupled pigments [10] and Förster theory for resonance energy transfer (FRET) for relatively weakly coupled pigments [11]. The effective light absorption and energy transfer properties of pigment-protein complexes make them desirable candidates for nanotechnological applications or at least a source of inspiration for synthetic designs [12, 13]. Bio-hybrid systems could utilise the natural properties of photosynthetic proteins in a useful device, for example, protein-sensitized solar cells often termed “biophotovoltaics” [14–18] and biosensors for the detection of herbicides [19, 20].

It is possible to augment LH proteins with other molecules in order to extend their absorption range so that light which falls within the aforementioned spectral gaps can be utilized. This is typically achieved by interfacing the protein with alternative pigments [21–24], or optically-active nanoparticles [25–27], which absorb strongly in complementary spectral regions where the natural absorption of the protein is minimal. Alternatively, complementary fluorescent proteins can be fused with natural LH proteins to enhance their absorption by using genetic modification strategies [28]. If these additional molecules are in close proximity to the protein and the spectral properties are favourable then energy will be absorbed and transferred to the protein’s natural pigments via FRET, effectively enhancing the absorption strength of the protein in the bio-hybrid system. The most common method of augmenting LH proteins is by direct covalent attachment of the donor molecule/nanoparticle [21–24, 27, 29]. A covalent linkage can be highly effective for ensuring a very short donor-acceptor distance but it does have some drawbacks, typically requiring either specific chemical reactions or time-consuming genetic modifications. Another limitation of many previous studies is that the augmented protein complexes were generally isolated in detergent micelles rather than their natural biomembrane location, which is a different physicochemical environment. A useful alternative system would be one that self-assembles, which does not require laborious chemical or genetic processes and that still provides enhanced light-harvesting capacity compared to natural systems. An ideal self-assembly approach: (i) would be applicable to any LH protein (e.g., both plant and bacterial), (ii) would allow a suite of different pigments to be used to fill the specific absorption ‘gaps’ for different proteins, (iii) would not need adaptation for different combinations of pigments and proteins, (iv) would utilize membranes. In this study, we investigate the use of a suite of membrane-localised dyes to enhance both plant light harvesting complex II (LHCII) and bacterial light-harvesting 2 complex (LH2) by co-reconstituting them into model lipid membranes formed via a simple self-assembly process. To test the effectiveness of this system, we address the following questions: (i) can different dyes be selected to enhance different spectral regions of various reconstituted LH proteins? (ii) can multiple dyes work in conjunction to fill more of the absorption gap of LH proteins? (iii) can dyes be added into pre-formed membranes after their initial assembly to allow multiple pigment additions? (iv) can the enhancement of LH proteins be demonstrated both for membranes suspended in solution and membranes deposited onto solid surfaces?

## 2. Materials and Methods

All chemicals were from Sigma-Aldrich (Merck), unless otherwise stated. Organic solvents were HPLC grade or higher and solid salts were BioUltra grade or higher. All water was deionized and passed through milli-Q purification system before use.

### 2.1 Lipids and dyes

All lipids and dyes were purchased in a dry, lyophilized form. The lipid-linked dyes (i) Texas Red 1,2-dihexadecanoyl-*sn*-glycero-3-phosphoethanolamine (TR-lipid), (ii) 1,2-dioleoyl-*sn*-glycero-3-phosphoethanolamine-ATTO647N (ATTO-lipid) and (iii) 1,2-dioleoyl-*sn*-glycero-3-phosphoethanolamine-Cyanine 7 (Cy7-lipid) were from Invitrogen, ATTO-TEC and Avanti Polar Lipids, respectively. The lipophilic dyes (i) 1,1’-dioctadecyl-3,3,3’,3’-tetramethylindocarbocyanine perchlorate (DiI) and (ii) 1,1’-Dioctadecyl-3,3,3’,3’-tetramethylindotricarbocyanine iodide (DiR) were both from Invitrogen. The standard lipids (i) 1,2-dioleoyl-*sn*-glycero-3-phospho-(1’-*rac*-glycerol) (DOPG) and (ii) soy asolectin lipid extract were from Avanti Polar Lipids and Sigma-Aldrich, respectively.

### 2.2 Biochemical purification of plant LHCII

Spinach leaves from a local supermarket were used as starting material. Trimeric LHCII complexes (the major antenna complex of photosystem II) were purified as described by Adams et al. [30] using the detergent *n*-dodecyl α-maltoside (α-DDM). SDS- and native-polyacrylamide gel electrophoresis (PAGE) and pigment extraction assays confirmed the protein’s purity and trimeric state (see Supplementary **Fig. S1**).

### 2.3 Biochemical purification of bacterial LH2

*Rba. sphaeroides* cells lacking the second LH2 operon (Δpuc2BA) and spheroidene monooxygenase (ΔcrtA) were grown in sealed 1 L glass Roux bottles under ~50 μMol m^-2^ s^-1^ for 72 hr [31]. This strain was selected to ensure a homogenous population of peptides and carotenoids were assembled into the LH2 complex. LH2 complexes were purified as described by Swainsbury et al. [32], except the detergent LDAO was exchanged for β-DDM at concentrations of 2% w/w for solubilisation and at 0.03% w/w, thereafter. The purified protein had an A_850_:A_280_ absorbance ratio of >3.0, confirming its high degree of purity.

### 2.4 Reconstitution of LH complexes and dyes into lipid vesicles

Glass vials and glass/steel Hamilton syringes were used whenever working with lipids or dyes in organic solvents. Dry lipids and lipid-linked dyes were dissolved into 2:1 chloroform:methanol and then mixed to generate the desired molar ratios and subsequently dried under nitrogen flow (40 min) and then placed in vacuum desiccator to remove any residual trace of organic solvents (10-16 hr, in the dark). The bulk lipid used for preparing samples for plant LHCII was soy asolectin lipid extract, whereas, the bulk lipid was DOPG for bacterial LH2. These lipids were chosen because they were previously established to provide a stable membrane environment for these specific protein complexes [33–35]. Aliquots of ~0.5 mg dry lipid mixture (as prepared above) were solubilised with an aqueous buffer of 0.5% α-DDM, 20 mM HEPES (pH 7.5) at room temperature for 12-16 hr with agitation to generate a mixed micellar lipid-DDM solution to a detergent-to-lipid molar ratio of ~9:1. A protein-lipid-detergent suspension was then prepared in plastic microfuge tubes by mixing calculated volumes of the following: lipid-DDM solution, aqueous buffers, and purified LH protein to produce a final concentration of 1 mM total lipid, 0.2% α-DDM, 20 mM HEPES (pH 7.5), 40 mM NaCl and the desired concentration of purified LH complexes. This protein-lipid-DDM suspension was then incubated with Bio-Beads SM-2 resin (Bio-Rad) to gradually remove the detergent (8, 20, 40 and 100 mg/mL for 1.5, 1.5, 1.5 and 16 hr, respectively). The lipids and membrane proteins self-assemble during the detergent removal process to form vesicles, termed proteoliposomes [36].

### 2.5 Spectroscopy of proteoliposomes

Before spectroscopy measurements, proteoliposome samples were diluted in a buffer of 40 mM NaCl, 20 mM HEPES (pH 7.5), to obtain a large enough volume for use in a 10 ×10 mm quartz cuvette (3 mL) at a low enough absorbance (~0.1 at 675 nm or 850 nm) to avoid inner filter effects [37]. Absorption spectroscopy was performed using an Agilent Technologies Cary 5000 UV-Vis-NIR absorption spectrophotometer. Fluorescence emission and excitation spectra were acquired immediately after absorption spectroscopy. During measurements, samples were maintained at 20 °C and gently stirred using a thermoelectrically cooled cuvette-holder with magnetic stirring capabilities (Quantum Northwest TC 1). Measurements on all LH2/dye and LHCII/TR samples were performed using an Edinburgh Instruments FLS980 fluorescence spectrophotometer equipped with a 450W Xenon arc lamp and dual excitation and emission monochromators. Scans for LH2/dye and LHCII/TR were collected using red-sensitive PMTs (Hamamatsu R928 or R980, respectively). Measurements on LHCII/DiI samples were performed using a Horiba Quantamaster fluorescence spectrometer equipped with similar specification Xenon arc lamp, dual monochromators and PMT detector. The two different fluorescence spectrometers were used simply due to equipment availability, therefore, to allow for fair comparisons, each dataset contained control samples of proteoliposomes without dyes, and all subsequent analysis was of “relative enhancement” levels as compared to these control samples. Data acquisition parameters were 0.5 nm step size, integrating 0.2 s/step and five scans averaged, for all measurements. Slit widths (spectral bandwidths) and wavelength range are specified in the main text figure captions. Spectra were analysed using Origin Pro (2019b) graphing software.

For assessing TR, Cy7 or ATTO, these lipid-linked dyes were included in the original lipid mixture (see section 2.4). Whereas, for assessing DiI/DiR, a solution of ~10.7 μM dye in ethanol and low volumes were injected into a pre-prepared sample of LHCII/LH2 proteoliposomes in a cuvette. Sequential absorbance and fluorescence spectra acquired ~15 min after each injection of dye solution.

### 2.6 Formation of hybrid thylakoid membranes on glass substrates

The polymerized lipid templates were prepared as described previously [38]. Briefly, lipid bilayers of 1,2-*bis* (10,12-tricosadiynoyl)-*sn*-glycero-3-phosphocholine (Diyne-PC) were deposited onto glass coverslip substrates by vesicle spreading and then polymerization was conducted by UV irradiation through a custom-designed photomask. Patterned substrates were stored in water, dried with nitrogen gas immediately before use, and placed into a microscopy sample holder. “Hybrid thylakoid membranes” were prepared as described by Meredith et al. [39]. Briefly, suspensions of thylakoid membranes (extracted from spinach) and lipid vesicles were combined in a 1:3 w/w ratio and incubated with a patterned glass substrate for 30 min. This generated supported membranes that were confined in 2-D into corrals. These samples were rinsed with copious buffer solution and then characterized by microscopy. TR-lipids and DiI dyes were incorporated into membranes as described in section 3.6.

### 2.7 Fluorescence lifetime imaging microscopy of hybrid thylakoid membranes

FLIM measurements were performed using a Microtime 200 time-resolved confocal fluorescence microscope (PicoQuant). This system uses an Olympus IX73 inverted optical microscope as a sample holder with light passing into and exiting various filter units for laser scanning, emission detection and timing electronics. The two excitation sources, 561 nm and 640 nm laser diodes, were driven in pulsed interleaved excitation mode by a PDL 828 Sepia II burst generator module (PicoQuant) at a pulse rate of 20 MHz to selectively excite either TR/DiI dye (561 nm) or the Chl within LH complexes (640 nm). The pulse width for the lasers were 70 ps and 90 ps, respectively. The laser power was set so that the excitation fluence at the sample was 0.026 mJ/cm^2^, found to be a good balance between achieving reasonable signal and avoiding annihilation effects, as previously reported [39]. The lasers were reflected toward the sample by a 561/640 (dual band) dichroic mirror with the beam focused through a 100× oil objective lens (N.A. 1.4) (UPlanSApo, Olympus). Emission from the sample was passed through the same objective and dichroic, towards the detectors. Emission light was separated by a 635LP beamsplitter so that emission <635 nm was directed though a 620/60 bandpass filter and to a hybrid PMT detector (TR/DiI emission) whilst emission >635 nm was directed though a 690/70 bandpass filter to a single-photon counting avalanche diode detector (Chl emission). Analysis of FLIM data was performed with SymPhoTime software (PicoQuant) and Origin Pro (2019b) graphing software.

## 3. Results and Discussion

### 3.1. Concept: testing multiple energy donors with two different LH complexes

Our previous work has shown that plant LHCII [40, 41] can be “enhanced” with lipid-linked Texas Red (TR) chromophores when they are co-assembled into model membranes in the form of either lipid vesicles [36] or lipid nanodiscs [42]. This TR moiety is amphiphilic and is tethered to lipid headgroups which allow the possibility of close contact with membrane proteins (**Fig. 1A(i)**) and, importantly, TR has absorption between 525-625 nm in the “green gap” of the LHCII absorption spectrum where there is minimal natural absorption (**Fig. 1C**). In these studies we demonstrated several advantages of using lipid-tagged chromophores: (i) the system is modular in the sense that dyes can be incorporated at a wide range of concentrations, (ii) LH proteins can be “enhanced” whilst in a biologically-relevant membrane environment, and (iii) membranes readily adsorb to surfaces making them amenable to surface-based nanotechnologies [36]. Evidence was presented for attractive interactions occurring between the TR chromophores and LHCII proteins which increased the rate of energy transfer [42]. For the current study, we hypothesized that it should be possible to enhance other LH complexes in a similar manner, and chose to study the LH2 antenna from *Rhodobacter* (*Rba.*) *sphaeroides* because it allows testing of an alternative spectral range compared to plant LHCII, with an absorption spectrum extending into the near-infrared. Furthermore, the *Rba. sphaeroides* LH2, referred to as LH2 henceforth, is a well-studied model protein for understanding light-harvesting in photosynthesis, giving a strong foundation for incorporating new pigments [32, 43–46]. LH2 has a spectral gap in the visible red region, between approx. 620-750 nm, and we chose two different chromophores which fit spectrally at different positions within this range: ATTO647N (hereafter, ATTO) and Cy7 (**Fig. 1D**). Both these dyes are commercially available conjugated to the headgroup of a DOPE lipid and were expected to co-assemble with LH2 during proteoliposome formation (**Fig. 1A(iv-vi)**), analogous to TR-lipids assembling with plant LHCII. Furthermore, we were interested in testing alternative methods for pigment incorporation that may have greater ease-of-use, so that they could be readily applied to other LH proteins of interest. We conceived that it might be possible to use relatively hydrophobic pigments that spontaneously localize to the hydrophobic interior of the lipid bilayer, and insert directly into pre-formed membranes. Therefore, we chose to test a series of dialkylcarbocyanine dyes which are commonly used as “lipophilic tracers” for whole-cell fluorescence microscopy [47]. These dialkylcarbocyanine dyes are dissolvable in polar organic solvents that are miscible with water, allowing them to be easily mixed with typical aqueous biological materials at any stage of sample preparation. These advantages allow for the simple injection of a few microliters of concentrated dialkylcarbocyanine in an ethanolic solution into an aqueous suspension of liposomes to deliver a significant quantity of exogenous pigment to the membrane (see **Methods 2.4**). The dialkylcarbocyanines DiI and DiR were chosen as potential energy donors to plant LHCII and LH2, respectively (**Fig. 1A(ii-iii)**), and for their appropriate spectral properties, as detailed in later sections.

**Figure 1.**
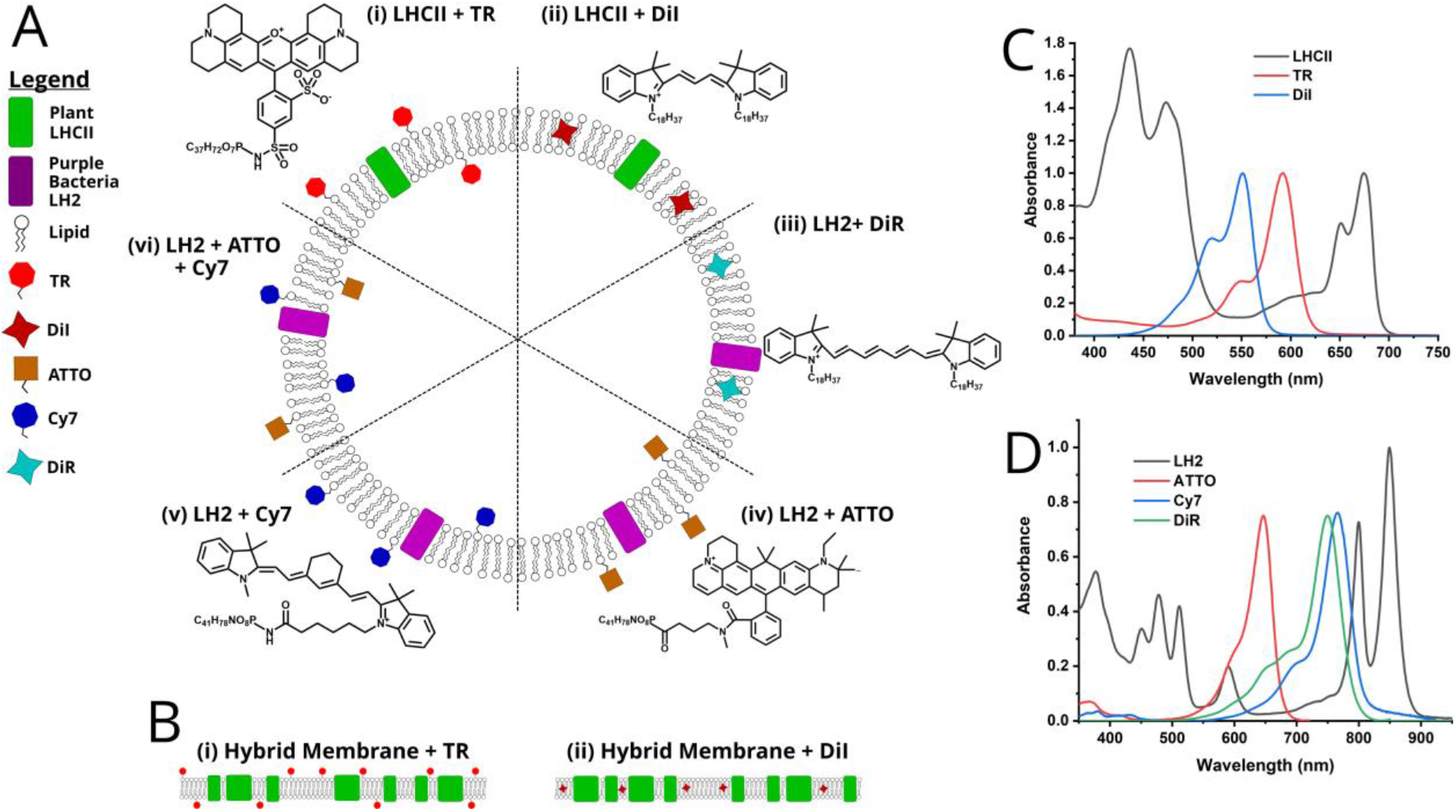
Schematic of the series of model membrane samples studied in this work. Proteoliposomes (**A**) were generated by combining lipids and biochemically-purified proteins in the desired molar ratios, following an established procedure that induces the spontaneous self-assembly of membrane vesicles (see **Methods** 2.4). When lipid-linked chromophores were used (TR, ATTO, Cy7) these were incorporated into the membrane during lipid and protein assembly, whereas the hydrophobic pigments (DiI, DiR) were injected into the buffer solution immediately prior to measurements, with multiple sequential injections possible. Full chemical structures of the dyes are given in the **Methods 2.1**. Supported membranes (**B**) were assembled onto hydrophilic glass coverslips, following a recently-developed procedure to generate “hybrid membranes” comprising a fusion of thylakoid membranes extracted from spinach chloroplasts and synthetic liposomes. Normalised absorption spectra of plant LHCII (**C**), and LH2 (**D**), with the additional dyes used to enhance the absorption of each protein. All dyes were obtained from commercial suppliers. The LH complexes were purified using established biochemical procedures (see **Methods 2.2+2.3** and **Fig. S1+S2**).

We can characterize the potential for energy transfer between proteins and extra pigments by assessing vesicular model membranes with solution spectroscopy, however, there is also interest in generating model membranes on solid support surfaces, because they are amenable to high-resolution microscopy [48–50] and could be compatible with surface-based device applications [51, 52]. Therefore, as a final test system we considered the possibility of enhancing our recently-reported “hybrid membranes” in which natural plant membranes are fused with synthetic membranes and assembled onto a planar glass surface [38, 39]. We considered that the LH complexes within these supported membranes could also be enhanced by incorporating either lipid-linked chromophores (TR) (**Fig. 1B(i)**) or free chromophores (DiI) (**Fig. 1B(ii)**) and we characterize these samples using fluorescence microscopy.

### 3.2. Expectations for each donor-acceptor pair from theory

We selected dyes that were spectrally well-matched to the proteins of interest, as described above, although each dye will have other photophysical properties that affect their capacity for FRET. It is necessary to assess how photophysical differences contribute to the effectiveness of each dye, and whether structural and chemical interactions between the dyes and the proteins modify these properties. One can expect the magnitude of the LH protein’s fluorescence enhancement (*F_LH_*) to be related to both the probability of photon absorption by the dye (*A_dye_*) and the efficiency of dye-to-protein FRET (*E_FRET_*), in other words: *F_LH_* ∝ *A_dye_* and *F_lh_* ∝ *E_FRET_*(*_dye→protein_*). The properties that absorption and FRET efficiency depend upon can then be considered. For the former:

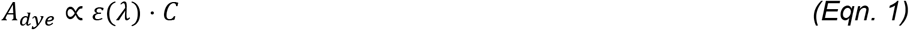

where *ε*(*λ*) is the dye’s molar absorption coefficient (a measure of its intrinsic absorbing strength), and *C* is the dye’s molar concentration. Then, assuming a certain number of dye molecules successfully promoted to an excited state, *E_FRET_* will depend upon the donor-acceptor separation distance (*r*) and their photophysical properties. Specifically, *E_FRET_* can be related to *r* by applying Förster theory [53]:

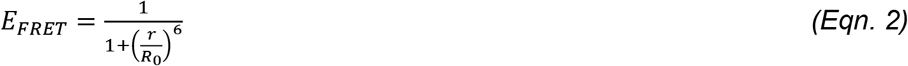

where *R_0_* is the distance at which the transfer efficiency is 50%, termed the Förster radius of the donor-acceptor pair. *R_0_* depends upon a combination of the photophysical properties of the system:

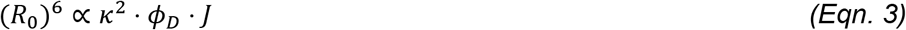

where, *κ^2^* represents the relative orientation between the donor and acceptor transition dipoles, *φ_D_* is the fluorescence quantum yield of the donor and *J* is the donor-acceptor spectral overlap integral. The full expressions for all properties discussed are detailed in the supplementary **Theory**.

From the above relationships, the effectiveness of each dye should be maximized by having: (i) broad absorption peaks and a high absorption coefficient, (ii) a high fluorescence quantum yield and (iii) a good spectral overlap with the LH protein. Values from the literature for the first two parameters are given for each dye in **Table 1**. In order to predict the effectiveness of the dyes as energy donors, the spectral overlap (*J*) and Förster radius (*R0*) were calculated for each dye-protein combination, see **Table 1**, supplementary **Theory 1**, and **Fig. S2**. A large value for both *R_0_* and *J* predicts a high FRET efficiency if the dyes are sufficiently close to the protein. Our experiments will measure the fluorescence enhancement of the protein and the FRET efficiency and these will be compared to the photophysical parameters discussed above. This allows us to determine whether the dyes have been properly incorporated into the lipid membranes and dispersed in a way that makes them structurally assemble to the protein.

**Table 1.** Photophysical properties collated from literature and Förster radius values calculated for each dye used in this study. “dye abs/em λ_max_”: wavelength of the main absorption/emission peak. ε: molar absorption coefficient. Φ: fluorescence quantum yield. J: spectral overlap integral. R_0_: Förster radius. Values for λ_max_, ε, Φ are taken from the reference listed. Values for J and R_0_ are calculated (see supplementary **Theory** text and **Fig. S2**). *Lipodots are dye-loaded oily droplets in an aqueous solution and the values for λ_max_, ε, Φ for DiI/DiR dyes in this solvent were used in place of values measured in aqueous solution (these dyes are not directly soluble in water).

| dye (donor) | LH protein (acceptor) | dye abs/em $\lambda_{max}$ (nm) | dye $\epsilon$ ($M^{-1} cm^{-1}$) | dye $\Phi$ | $J$ ($M^{-1} cm^{-1} nm^4$) | $R_0$ (Å) | solvent | reference |
| --- | --- | --- | --- | --- | --- | --- | --- | --- |
| TR | plant LHCII | 589/ 610 | 116,000 | 0.35 | $1.03 \times 10^{17}$ | 88 | aqueous | [54] |
| Dil | plant LHCII | 549/ 790 | 144,000 | 0.31 | $5.93 \times 10^{16}$ | 79 | Lipiodot* | [55] |
| ATTO | LH2 | 646/ 664 | 150,000 | 0.65 | $4.36 \times 10^{16}$ | 85 | aqueous | [56] |
| Cy7 | LH2 | 747/ 779 | 250,000 | 0.28 | $4.54 \times 10^{17}$ | 109 | aqueous | [57] |
| DiR | LH2 | 748/ 780 | 270,000 | 0.21 | $3.74 \times 10^{17}$ | 101 | Lipiodot* [55] | |

### 3.3. Comparison of the effectiveness of TR and DiI for increasing the spectral range of plant LHCII

Previously [36], we showed that plant LHCII in liposomes could be enhanced with TR but only considered a limited range of chromophore concentrations up to a TR:LHCII molar ratio of ~20:1. Surprisingly, the trend for the enhancement of LHCII fluorescence appeared to be linear, so we speculated that the enhancement must saturate well in excess of a 20:1 ratio. In the current work, we assess the absolute limit of LHCII enhancement by TR, by extending the ratios up to ~80:1. A sample set of LHCII-TR proteoliposomes was prepared, starting with a constant concentration of the purified LHCII (~0.6 μM) and lipid (1 mM) and a series of increasing TR concentrations (9-45 μM). Absorption spectra showed that sample preparation had been successful, whereby the LHCII Chl *a* Q_y_ transition at ~675 nm was relatively consistent between samples and the TR peak at 590 nm with a vibronic shoulder at 550 nm increased linearly with the dye concentration (**Fig. 2A**). For all samples, the actual dye-to-protein ratio achieved was quantified by analysing the absorbance spectra of the assembled liposomes (see Supplementary **Table S1**). Fluorescence emission scans were acquired by selectively exciting the dye at 540 nm (where LHCII absorption is very low) and collecting broad-range emission spectra to distinguish the fluorescence emission from the dye (direct re-emission) or from the LH2 protein (FRET then emission). These fluorescence emission spectra showed strong enhancement of the LHCII emission peak at ~680 nm, increasing with TR concentration to more than 3× the fluorescence of LHCII alone (**Fig. 2B**). This is good evidence for FRET from TR to LHCII, as observed in previous work [36]. Note that the magnitude of the TR emission peak at ~610 nm does not increase proportionally with TR concentration (**Fig. 2E**)suggesting that it becomes self-quenched at high concentrations (supplementary **Fig. S3** and **Table S2**) [58].

**Figure 2.**
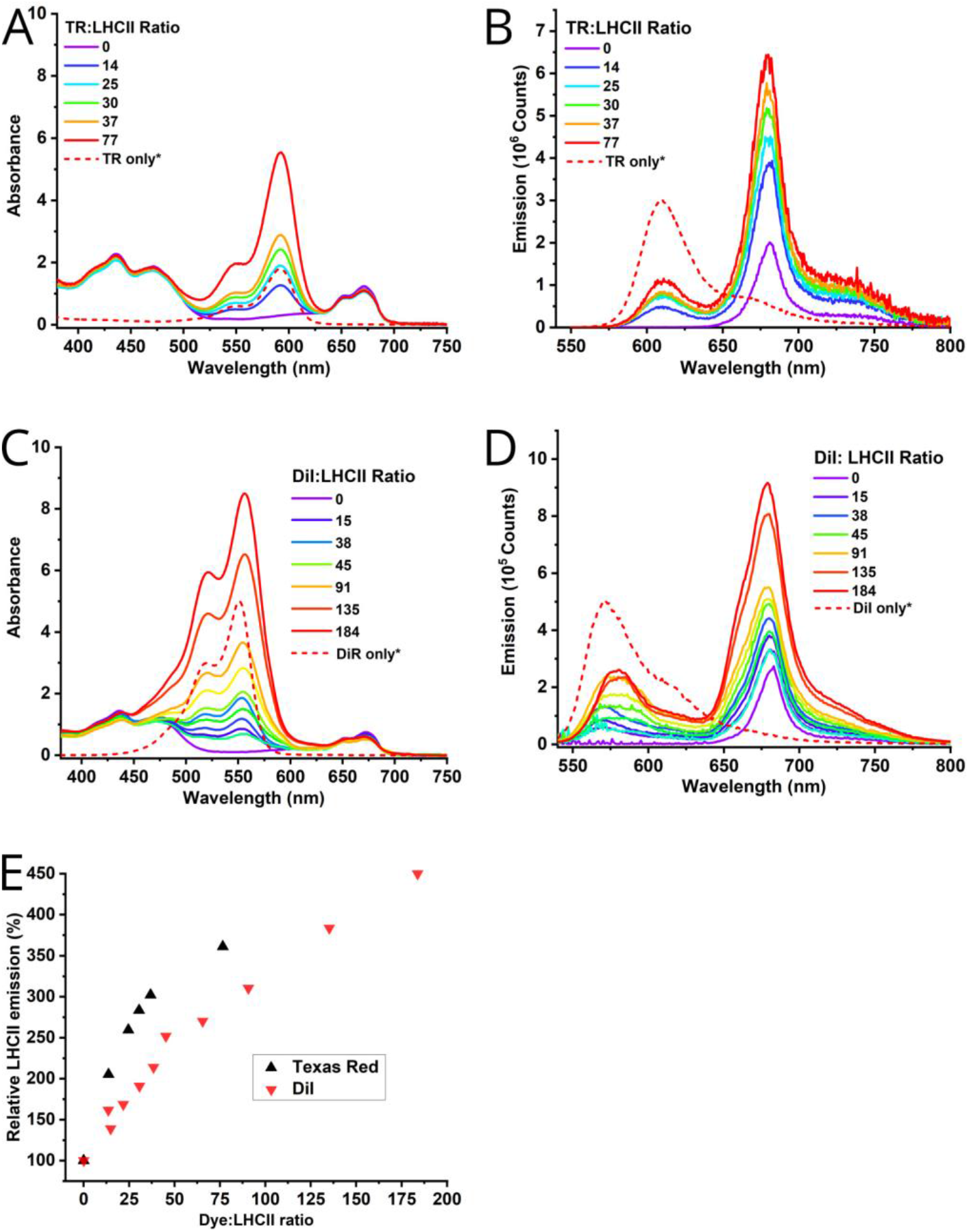
Comparison of the effectiveness of two different types of chromophore for enhancing plant LHCII. (**A**) Absorption spectra of LHCII proteoliposomes containing lipid-linked TR at the concentrations shown. *Indicates normalised reference data. (**B**) Fluorescence emission spectra of the same samples as in (A) with selective excitation of TR at 540 nm, ex/em bandwidths 1/1nm. (**C**) Absorption spectra of LHCII proteoliposomes after injection of DiI to the concentrations shown. (**D**) Fluorescence emission spectra of the same samples as in (C), with selective excitation of DiI at 525 nm, em/ex bandwidths 2/2 nm. Selective excitation wavelengths for dyes were chosen to minimise direct LHCII excitation rather than to be centred at their peak excitation wavelengths. Legend for each spectrum not shown for visual clarity; the full table of results are shown in supplementary **Tables S1** and **S3**. (**E**) Graph comparing the relative enhancement effect between these dyes, where 100% is the intensity of fluorescence at 680 nm in LHCII proteoliposomes in the absence of any dye. The LHCII emission intensity was calculated after subtraction of the overlapping fluorescence due to the dye, the full calculations are shown in supplementary **Table S3**.

Next, the idea of assembling dialkylcarbocyanine dyes from solution into pre-formed proteoliposomes was tested. To do this, the DiI dye was sequentially injected into a sample of LHCII proteoliposomes and spectra were acquired after each addition. This was repeated to cover a series of different ranges of DiI concentration. Absorption spectra of the resulting LHCII-DiI proteoliposomes (**Fig. 2C**) show similar height peaks for LHCII at ~675 nm and the expected range of DiI content is evident from the sequence of double peaks at ~560/520 nm. A greater final concentration of DiI was easily achieved with this injection method, up to a measured DiI:LHCII ratio of 184:1, as compared to the more time-consuming co-assembly method required when utilizing lipid-linked TR (77:1). Fluorescence emission spectra (**Fig. 2D**) showed an enhancement of the main (681 nm) LHCII fluorescence peak up to nearly 4× its original level. A shoulder at ~660 nm is apparent on the LHCII peak at the higher TR:LHCII ratios due to the contribution from the vibronic tail of the TR emission. Unlike for TR-LHCII, DiI produces a relatively linear increase in LHCII fluorescence intensity relative to its concentration suggesting that dye self-quenching is not significant at these concentrations.

The effectiveness of these two dyes was compared by plotting the enhancement of LHCII fluorescence against the dye:protein ratio for each (**Fig. 2E**). The TR dye appears to generate higher levels of LHCII fluorescence than DiI at similar dye concentrations, i.e., TR appears to be a more effective donor of excitation energy to LHCII. This is supported by calculations of energy transfer efficiency (ETE), determined by a comparison of the fluorescence excitation and linear absorption spectra of selected samples (see Supplementary **Table S4**), which found that ETE is between 60-97% for TR-LHCII and only 25-33% for DiI-LHCII. The significantly higher FRET efficiency must be due to a stronger average coupling between TR+LHCII than between DiI+LHCII and could be due either to the difference in spectral overlap (i.e., energetic coupling) and/or due to different distributions and thus separation distances between protein and dye molecules (i.e., dipole-dipole coupling). For TR-LHCII and DiI-LHCII interactions, the values for *J* (spectral overlap) were calculated as 1.03×10^17^ and 5.93×10^16^ M^-1^ cm^-1^ nm^4^, respectively, and *R*_0_ (Förster radius) as 88 Å and 79 Å, respectively, as shown in **Table 1**. This supports the possibility that spectral overlap is a major reason why the observed ETE is higher for TR than DiI. It could also be that the TR-lipids localize at shorter distances to LHCII than DiI does within the membrane. The maximum enhancement observed was ~361% for TR at a dye:protein ratio of 77:1 whereas enhancement due to DiI continued to increase with an apparently linear trend up to ~450% at a dye:protein ratio of 184:1. It was not possible to generate stable proteoliposomes with >80:1 TR/protein or >180:1 DiI/protein presumably due to lipid-lipid assembly issues, which could arise because both TR and DiI are bulky; furthermore, TR is charged and unfavourable repulsive interactions may occur. The enhancement effect when using TR but not DiI was also limited by fluorescence self-quenching (the propensity of different molecules to self-quench is not trivial to define). This meant that although DiI had a lower overall ETE due poorer energetic coupling it induced a greater absolute level of LHCII enhancement than TR because higher DiI concentrations could be assembled into membranes. Altogether, these findings show that different structural, physicochemical and spectral properties may alter the effectiveness of any given dye, although these may be difficult to distinguish. This will be explored further in the next section.

### 3.4. Comparison of the effectiveness of three different dyes for increasing the spectral range of LH2

Whilst other studies have assessed how covalent attachment of synthetic pigments to the LH2 polypeptides could enhance the absorption range of LH2 [59, 60], the effectiveness of self-assembled dyes for enhancing this protein has not been explored. To do this, first, we studied a series of proteoliposome samples with similar LH2 concentrations assembled with a range of concentrations of Cy7-lipids. The absorption of Cy7 appears at ~770 nm overlapping with the B800 LH2 BChl peak at 800 nm, and is observed as a shoulder on the blue-side of the B800 band at low Cy7 concentrations, becoming dominant at higher Cy7 concentrations (**Fig. 3A**). The dye-to-protein ratio and the dye incorporation yield for all LH2 proteoliposomes was quantified (see supplementary **Table S1**). Fluorescence emission spectra acquired with selective excitation of the Cy7 chromophore show an increasing intensity for both the Cy7 (790 nm) and LH2 (860 nm) peaks as the concentration of Cy7 is increased across the sample range, which is good evidence of FRET from Cy7 to LH2 (**Fig. 3B**). The emission spectrum of Cy7 has excellent energetic overlap with the B800 BChl absorption of LH2 (J ≈ 4.5×10^17^ M^-1^cm^-1^nm^4^, **Table 1**) so effective transfer would be expected, assuming that the dye is in close proximity to the protein. To test the effect of different spectral overlaps, ATTO-lipids were incorporated as a lipid-linked dye which absorbs over an alternative range, filling a different part of the spectral gap of LH2 and having a much weaker energetic overlap with the B800 BChl Qy transition (J ≈ 4.4×10^16^ M^-1^cm^-1^nm^4^). The absorption of ATTO appears at ~650 nm and has slight overlap with the LH2 peaks representing the BChl Q_X_ transitions at ~590 nm, and with the B800 absorption band (**Fig. 3C**). Emission spectra acquired for this sample set with selective excitation of the ATTO were quite weak compared with the emission from ATTO, although LH2 fluorescence is still greatly enhanced compared to the level in the absence of any synthetic chromophores at this excitation wavelength (**Fig. 3D**). When the two different lipid-linked pigments are mixed and incorporated into LH2 proteoliposomes a combinatorial effect is observed and the enhancement of LH2 fluorescence is greater than using either dye alone (**Fig. 3E-F**). The effect of using two dyes together can be quantified more easily with alternative measurements that assess a range of excitation wavelengths, presented later. Next, a dialkylcarbocyanine dye that is spectrally-compatible with LH2 was trialled. DiR was injected into pre-formed LH2 proteoliposomes. The absorbance spectra show a broad and inhomogeneous band between 600-750 nm representing DiR at increasing concentrations (**Fig. 3G**), which could suggest that DiR was unstable in liposomes and had aggregated. The calculated incorporation yield of DiR into LH2 proteoliposomes was relatively low (10-15% of the injected DiR was actually detected in the final sample, see supplementary **Table S1**), meaning that a relatively low dye:LH2 ratio of ~20:1 was achieved even after injecting large quantities. Nevertheless, emission spectra showed that there was significant enhancement of LH2 fluorescence at the highest DiR concentrations (**Fig. 3H**).

**Figure 3.**
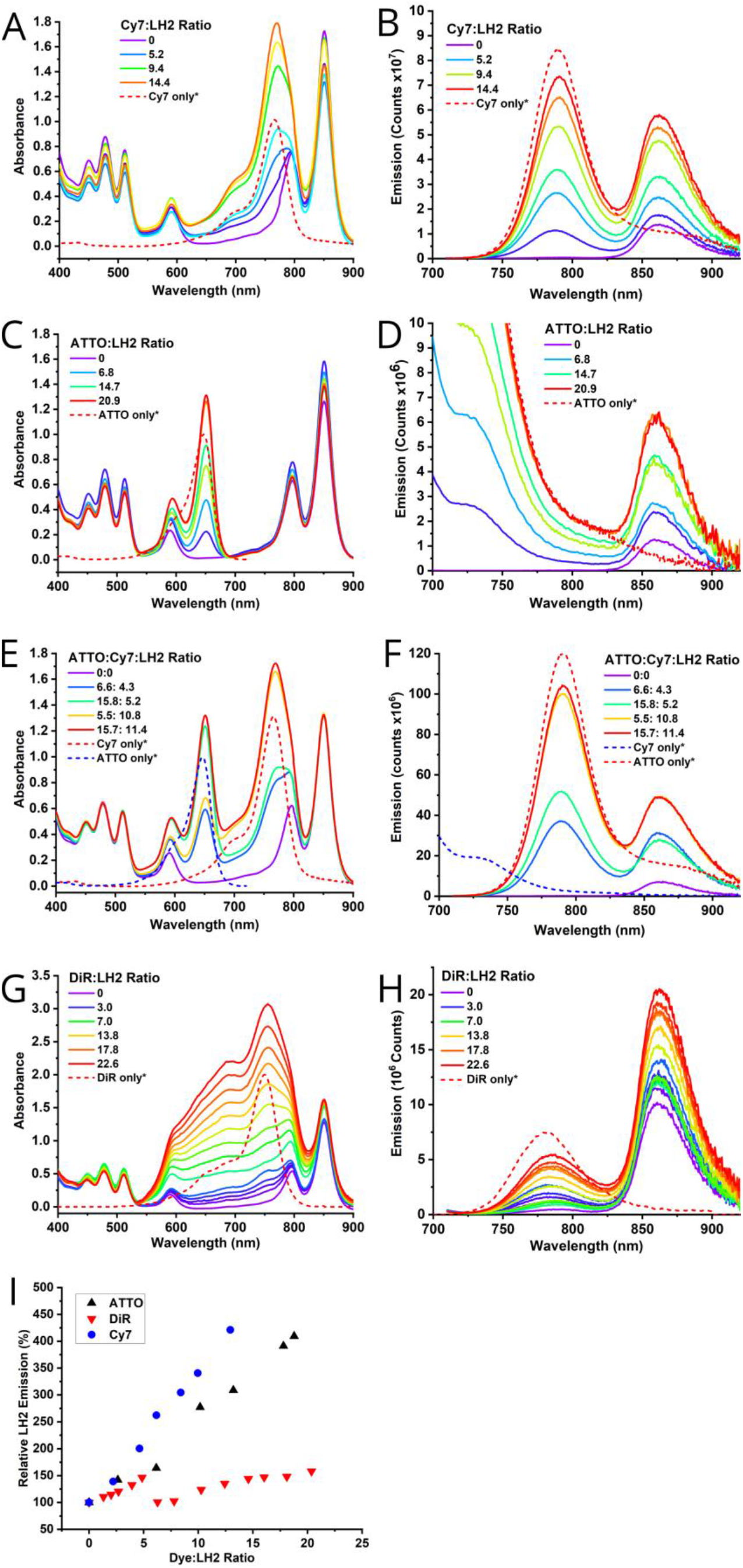
Comparison of the effectiveness of three different types of chromophore for enhancing fluorescence emission from the LH2 complex. Absorption spectra of LH2 proteoliposomes containing either lipid-linked Cy7 (**A**), lipid-linked ATTO (**C**), both Cy7 and ATTO lipid-linked dyes together (**E**), at the concentrations shown. *lndicates normalised reference data. Fluorescence emission spectra of the matched samples is shown, with selective excitation of Cy7 at 700 nm, ex/em bandwidths 4/4 nm (**B),(F**) or ATTO at 625 nm, ex/em bandwidths 2/2 nm (**D**). Absorption spectra of LH2 proteoliposomes after injection of DiR in ethanolic solution (**G**) to the concentrations shown. (**H**) Fluorescence emission spectra of the same samples as in (G), with selective excitation of DiR at 700 nm, em/ex bandwidths 4/4 nm. Selective excitation wavelengths for dyes were chosen to minimise direct LH2 excitation rather than to be centred at their peak excitation wavelengths. Legend for each spectrum not shown for visual clarity; the full table of results are shown in supplementary **Tables S1** and **S2**. (**I**) Graph comparing the relative enhancement effect between these dyes, where 100% is the intensity of fluorescence at 865 nm in LH2 proteoliposomes in the absence of any dye. The LHCII emission intensity was calculated after subtraction of the overlapping fluorescence due to the dye, the full calculations are shown in supplementary **Table S3**.

The effectiveness of these dyes in terms of their enhancement of the fluorescence intensity of LH2 was compared against the dye:protein ratio (see **Fig. 3I**). Note, this analysis compares the relative change in fluorescence intensity of LH2, due to the presence of donors as compared to their absence, when using an idealized wavelength that selects for excitation of the particular dye. Cy7 and ATTO are both effective at enhancing LH2 fluorescence emission with Cy7 appearing to be slightly superior, maximum enhancements of 421% and 409% are observed at dye/protein rations of 12.9: 1 and 18.8: 1 for Cy7 and ATTO, respectively (*blue* vs. *black datapoints*, **Fig. 3I**). This may be explained when one considers both the spectral overlap relevant for FRET and the background level of LH2 absorbance at different wavelengths. First, we note that the spectral overlap integral between LH2 and the dye is significantly lower for ATTO-to-B800 than Cy7-to-B800 and this is reflected in the values calculated for Förster radii of 85 Å and 109 Å for ATTO-LH2 and Cy7-LH2, respectively (see **Table 1**). This correlates with a lower average ETE calculated from linear absorption versus excitation spectra of 53-66% for ATTO-LH2 compared to 60-91% for Cy7-LH2 (see supplementary **Fig. S4** and **Table S4**). However, the absorption of the LH2 complex is slightly lower at 625 nm compared to 700 nm, the wavelengths used in these experiments to selectively excite either ATTO or Cy7, respectively, and this subtle difference actually leads to a relatively large enhancement factor for ATTO with 625 nm excitation light. Conversely, DiR appears to be much less effective as an excitation donor to LH2 than either Cy7 or ATTO at the similar concentrations with a maximum enhancemnt of 158% at a dye: protein ratio of 20.4: 1, less than 1/5 of the level with ATTO and Cy7 (*red datapoints*,**Fig. 3I**). This correlates with the relatively low ETE calculated for DiR to LH2 in these samples with a range of 3-11% (see supplementary **Fig. S4** and **Table S4**). Considering that the spectral overlap for DiR-LH2 is very similar to Cy7-LH2, leading to a similar Förster radius of 101 Å vs. 109 Å (**Table 1**), then this implies that the much lower measured ETE is due to less favourable structural interactions between DiR and the protein. In other words, the average distance between DiR and LH2 must be larger than between Cy7-lipids and LH2 even for the same ratio of dye:LH2. The most reasonable explanation is that some physicochemical factors limit the molecular interactions between DiR and LH2, for example, repulsive electrostatic interactions between DiR and the protein or due to aggregation of DiR. The latter possibility is supported by broadening of the absorbance peaks of DiR at high dye:protein ratios (see **Fig. 3G**). Overall, our findings on the potential for enhancing LH2 fluorescence with dyes show that spectral overlap is very important but that favourable structural interactions allowing proper dye incorporation into lipid bilayers are a prerequisite.

### 3.5. Assessment of the overall enhancement of LH complexes by consideration of the increased peak area in fluorescence excitation spectra

In the previous sections, we assessed the dye-to-protein ratio from absorbance spectra and the “enhancement” of the LH protein’s fluorescence by monitoring the relative increase in the intensity of its emission peak. It is also informative to assess the overall performance of the system over a broad range of illumination wavelengths where the synthetic dyes may or may not be excited, rather than only selecting the ideal wavelength for dye excitation. To do this, fluorescence excitation scans were acquired on LH2 proteoliposomes. Here the detection wavelength was fixed at a position that only monitors fluorescence emission from LH2 (886 nm) and a full range of excitation wavelengths was scanned (450-875 nm). The magnitude and breadth of these additional peaks relative to the peaks attributed to the pigments bound to LH2 represents the effective increase of the absorption range. We use the term “effective increase in absorption” because only those absorption events which lead to FRET and subsequent emission from LH2 appear as a signal here. Fluorescence excitation spectra for LH2 proteoliposomes are shown side-by-side for all dyes in **Fig 4** and the enhancement effect can be quantified as the area under these peaks. Firstly, comparing data for the lipid-linked dyes (**Fig. 4A-B**), it is apparent that Cy7 produced significantly greater enhancement than ATTO, although the latter was still effective. For example, at ~5:1 ratio of Cy7:LH2 the effective increase in absorption area related to the dye between 600-800 nm was similar to the original area of the LH2 B800 peak (*blue spectrum*,**Fig. 4A**) and at ~14:1 Cy7:LH2 the increase in area was greater than the LH2 B850 peak (*red spectrum*,**Fig. 4A**). Whereas, at the maximum-tested ratio of ~21:1 ATTO:LH2 the new band at ~650 nm only reached an area similar to the LH2 B800 transition (*red spectrum*,**Fig. 4B**). In comparison, the enhancement of LH2 achieved with DiR using the injection method was smaller, where the peak related to the dye at ~750 nm was more subtle, reaching an area of approximately half of the LH2 B800 band at the highest ratio of ~23:1 DiR:LH2 (*red spectrum*,**Fig. 4C**). The combination of both Cy7 and ATTO led to the most impressive results with prominent peaks at both ~650 nm and 770 nm, varying in magnitude according to the ratio of the two dyes that were included (**Fig. 4D**). At the highest ratio of ~16:11:1 ATTO:Cy7:LH2 the enhancement effect was large enough to approximately double the original absorption area of LH2 considering the full range 450-875 nm (*red* vs. *purple spectra*,**Fig. 4D**). The process of analysing fluorescence excitation spectra was also performed for plant LHCII proteoliposomes and it was clear that both the TR and DiI dyes produced sufficient enhancement in the green spectral region, 525-625 nm, significantly enhancing the absorption of the complex across the full range (see supplementary **Table S5**).

**Figure 4.**
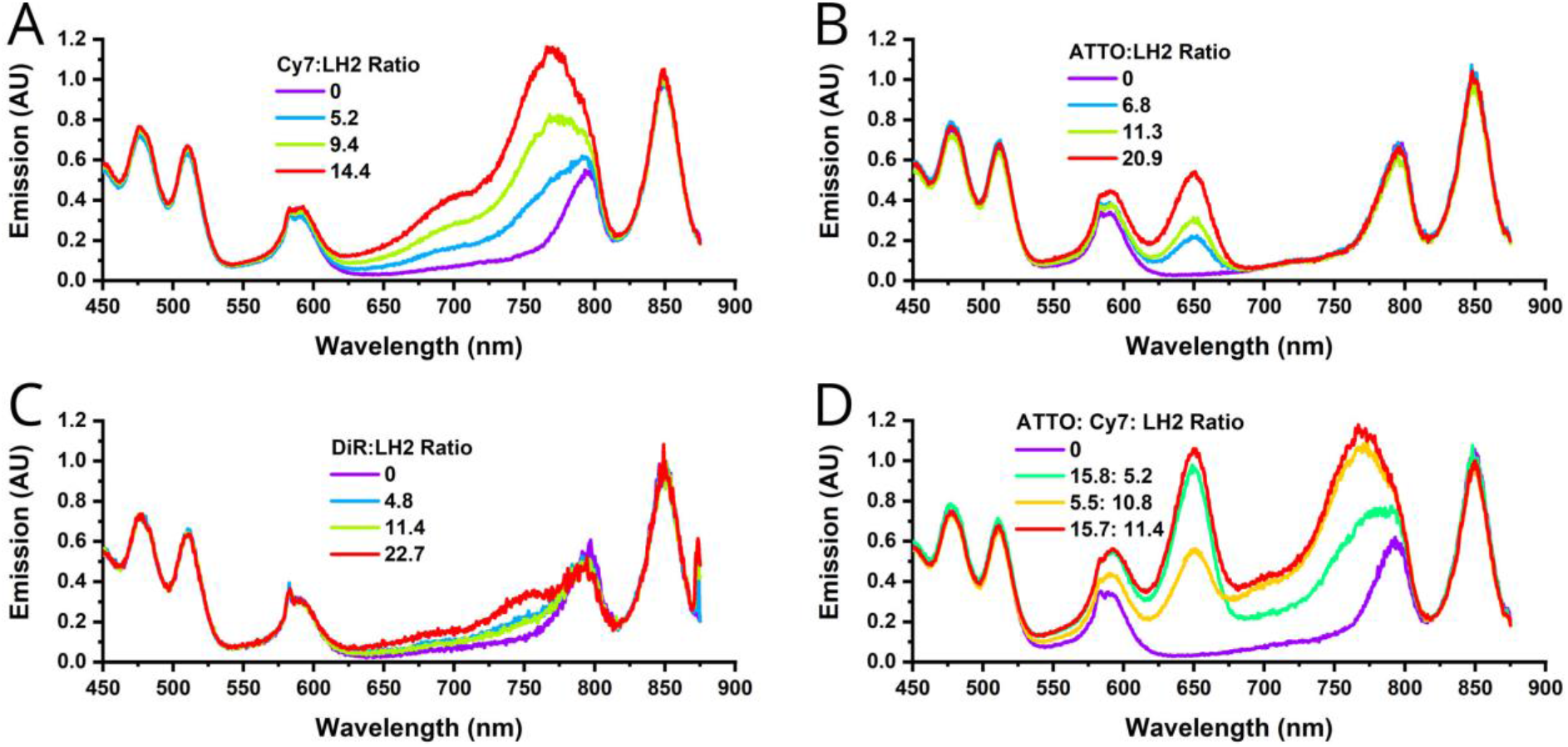
Comparison of the relative enhancement of LH2 by assessment of fluorescence excitation spectra. All spectra were acquired with the selective detection of LH2 emission at 875 nm, ex/em bandwidth 4/2 nm. Fluorescence excitation spectra for LH2 proteoliposomes containing either Cy7-lipids (**A**), ATTO-lipids (**B**), injected DiR (**C**), or both Cy7-lipids and ATTO-lipids together (**D**). Fluorescence excitation spectra for plant LHCII samples are shown in supplementary **Fig. S5**.

To quantify how each dye enhances LH2 and LHCII complexes, the fluorescence excitation data were analysed in more detail. The effective enhancement was calculated as the increase in the integrated area under each spectrum, normalized so that an area of 100% represents the natural absorption strength of the particular LH protein across the full measurement range. This highlights the additional absorption due to the presence of the dye. Comparison between plant LHCII and LH2 enhancement was achieved by converting the dye-to-protein ratio to a dye-to-(B)Chl ratio (mole/mole) for each sample by accounting for the number of Chl-type pigments known to be present in each LH complex (42 Chls in LHCII, 27 BChls in LH2). This analysis is shown for all dye/protein combinations in **Fig 5**. Two general comments can be made. Firstly, it was possible to increase the absorption both LH proteins by close to 100% (i.e., to 200% of the natural absorption strength), but less dye was required for LH2 than plant LHCII (~1 dye:BChl versus >2 dye:Chl) to achieve the same level of enhancement. Secondly, the trends for effective absorption appear to plateau only for the very highest dye-to-Chl ratios reached for TR lipids and, surprisingly, the maximum enhancements for LH2 with Cy7 and Cy7+ATTO have not yet been reached, nor for LHCII with DiI. Note that this analysis considers the absorption across the full spectral range as a fair way to place the enhancement effect in the context of an entire LH protein complex exposed to unfiltered white light. If we choose to focus only on the absorption in “spectral gap” regions then the effective absorption compared to the very low natural levels reaches a maximum of 489% for plant LHCII with DiI and 465% for LH2 with combined ATTO and Cy7 (see supplementary **Table S5**).

**Figure 5.**
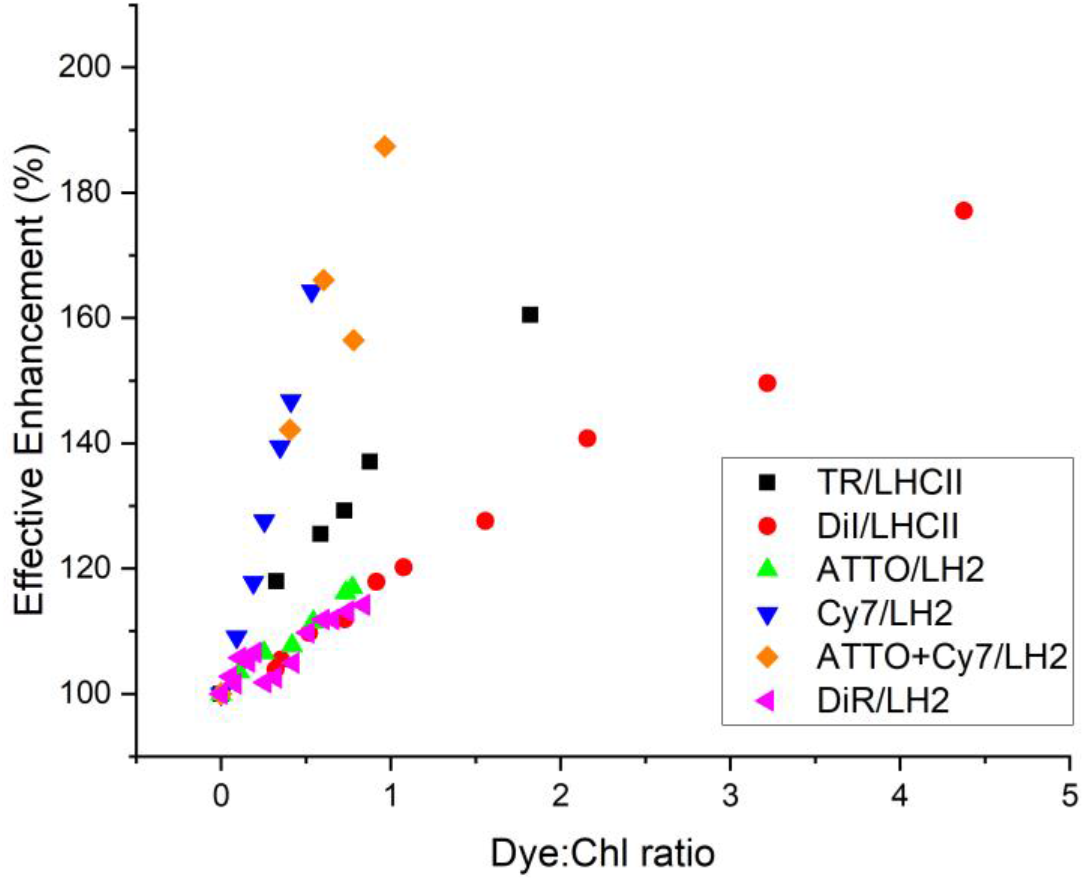
Comparison of LH protein enhancement between all dye and protein combinations, considering an extended spectral range. The “effective absorption” was calculated from the integrated areas under the fluorescence excitation spectra for LH2 (**Fig. 4**) and plant LHCII (**Fig. S2**), and normalizing so that 100% absorption (380-680 nm and 450-875 nm for LHCII and LH2, respectively) represents the natural (unmodified) protein complex. The numerical data are tabulated in see supplementary **Table S5**. The molar Dye:Chl ratio was calculated as [dye:protein]×42 for plant LHCII and [dye:protein]×27 for plant LHCII, using the dye:protein ratios already determined from absorption spectra (see supplementary **Table S1**).

Beyond these general trends, we can gain further insight by considering the differences between types of dye, as explained below. At the highest concentrations of TR and DiI, the effective absorption of plant LHCII was enhanced to 160 and 177%, respectively. This follows a trend of increasing enhancement with increasing dye content and shows that TR is slightly more effective at lower concentrations compared to DiI (**Fig 5**, *black* vs *red datapoints*). This reinforces the finding that the simple method of injection and spontaneous insertion of lipophilic dyes into pre-formed proteoliposomes has the potential to enhance an LH complex as strongly as a lipid-linked dye, although greater quantities of the lipophilic dye may be required. For LH2, the effective absorption strength increased more rapidly with Cy7 than ATTO and reached a greater absolute level of 164% vs. 117% at the highest dye concentrations measured (**Fig 5**, *blue* vs *green datapoints*). This could be partially attributed to the much greater spectral overlap of Cy7 with LH2 than ATTO with LH2, as noted earlier. Furthermore, the ATTO dye has a lower molar absorption coefficient and a slightly narrower absorption peak than Cy7, which limits its absorption strength and range (ε_646 nm_[ATTO] = 150,000 M^-1^ cm^-1^; ε_747nm_[Cy7] = 250,000 M^-1^ cm^-1^). We cannot rule out the possibility that the Cy7-lipids simply have more favourable packing with LH2 than the ATTO-lipids, due to different chemical interactions, and because of the sixth-power relationship between donor-acceptor separation distance and transfer rate even small variations in average distance will have major effects on the level of enhancement. The highest absolute enhancement of LH2 was achieved by the combination of high levels of ATTO and Cy7 to a maximum of 194% (**Fig 5**, *orange datapoints*). This may be further explained by the finding that energy absorbed by the ATTO has two potential routes towards LH2, either directly (ATTO→ LH2) or via Cy7 (ATTO→ Cy7→ LH2) (data in supplementary **Fig. S6** and **Table S6** confirms that ATTO→ Cy7 FRET occurs). DiR provided the least enhancement of LH2 to an effective absorption of just 114% (**Fig 5**, *pink datapoints*). Whilst the trends over the low-concentration range for DiR-LH2 and DiI-LHCII were similar, overall, DiR may be considered much less effective because higher dye:protein ratios simply could not be achieved (due to the low incorporation yield of DiR noted earlier).

Overall, we can highlight the factors that dominate the ability of a dye to enhance the absorption strength of LH complexes by a final comparison of the predicted potential for FRET from theory (**Table 1**) to the experimental data from actual assembled membranes (**Fig. 5**): (i) a strong spectral overlap resulted in a high FRET efficiency and explained why TR was more effective than DiI and why Cy7 more effective than ATTO, (ii) a larger bandwidth of absorption peaks and absorption coefficient was also beneficial (Cy7 > ATTO), (iii) the differences in fluorescence quantum yield did not appear to be significant (higher *Φ* for ATTO did not rescue its performance), (iv) the ability to assemble high dye-to-protein ratios was crucial for overall effectiveness and presumably resulted in high packing densities of dyes in the membrane (DiI >> DiR). Thus, the dicarboxylate dye DiI was found to be an effective alternative to lipid-linked TR, despite a lower FRET efficiency, because very high dye:protein ratios could be accessed with the injection method (and self-quenching did not occur). Whereas, the other dicarboxylate, DiR, could not compete with lipid-linked Cy7 due to both poor incorporation yield and a low FRET efficiency. These findings show the importance of investigating different dyes and highlight the fact that physicochemical factors are critical in the self-assembly processes.

### 3.6. Assessing the potential for enhancing surface-associated membranes

Finally, we wished to assess the feasibility of using dyes in surface-supported membranes. Recently, we reported a new model system of bio/hybrid photosynthetic membranes that are amenable to high-resolution surface-based microscopy. These “hybrid thylakoid membranes” are generated from a combination of natural membranes extracted from spinach chloroplasts and synthetic lipid vesicles that fuse on a glass surface to form large-scale array patterns [39]. These surface-supported membranes are confined to corrals of 20 × 20 μm by using a pre-designed template (see **Methods** section 2.6). In order to test whether our strategy of self-assembly of donor chromophores into lipid bilayers could be effective for enhancing LH complexes within supported membranes, experiments were performed where hybrid membranes were supplemented with additional dyes. Both of the dye incorporation methods studied earlier in this paper were trialled, i.e., inclusion of TR-lipids at the lipid preparation stage or DiI injection after membrane formation. Fluorescence Lifetime Imaging Microscopy (FLIM) was performed to quantify both the donor quenching and acceptor enhancement which would be indicative of FRET. The FLIM instrument was set up to generate three separate image channels: (i) the *“Dye channel”* had selective TR/DiI excitation (561 nm) and selective collection of TR/DiI emission (590-635 nm), (ii) the *“Chl channel”* had selective excitation of the Chl within LH proteins (640 nm) and selective collection of Chl emission (590-635 nm), and (iii) *“Chl enhancement”* channel had selective TR/DiI excitation (561 nm) and selective collection of Chl emission (590-635 nm). To test TR, three different types of sample were compared: (i) hybrid membranes, (ii) lipid membranes containing TR-lipids, (iii) hybrid membranes including TR-lipids (*top*, *middle*, or *bottom* row of panels in **Fig. 6A**). Considering the *TR channel*, the fluorescence intensity was somewhat lower in the combined sample (**Fig. 6A(vii)**) as compared to the TR-only sample (**Fig. 6A(iv)**). Concurrently, the fluorescence lifetime of TR was significantly reduced between these samples, a strong indication of TR-to-Chl FRET (*green* vs. *yellow* colour, equating to a mean lifetime of 1.9 ns vs. 3.7 ns). Considering the *Chl channel*, the intensity was similar in the hybrid membrane samples irrespective of the presence of TR (**Fig. 6A(ii)** and (**viii**)), as expected. In contrast, for the *Chl enhancement* channel representing Chl-protein fluorescence after selective excitation of TR, there was much greater signal in the hybrid membranes containing TR as compared to the control samples (**Fig. 6A(ix)** compared to either (**iii**) or (**vi**)). This is good evidence for FRET from TR dyes to either LHCII or photosystem complexes within these mixed membranes.

**Figure 6.**
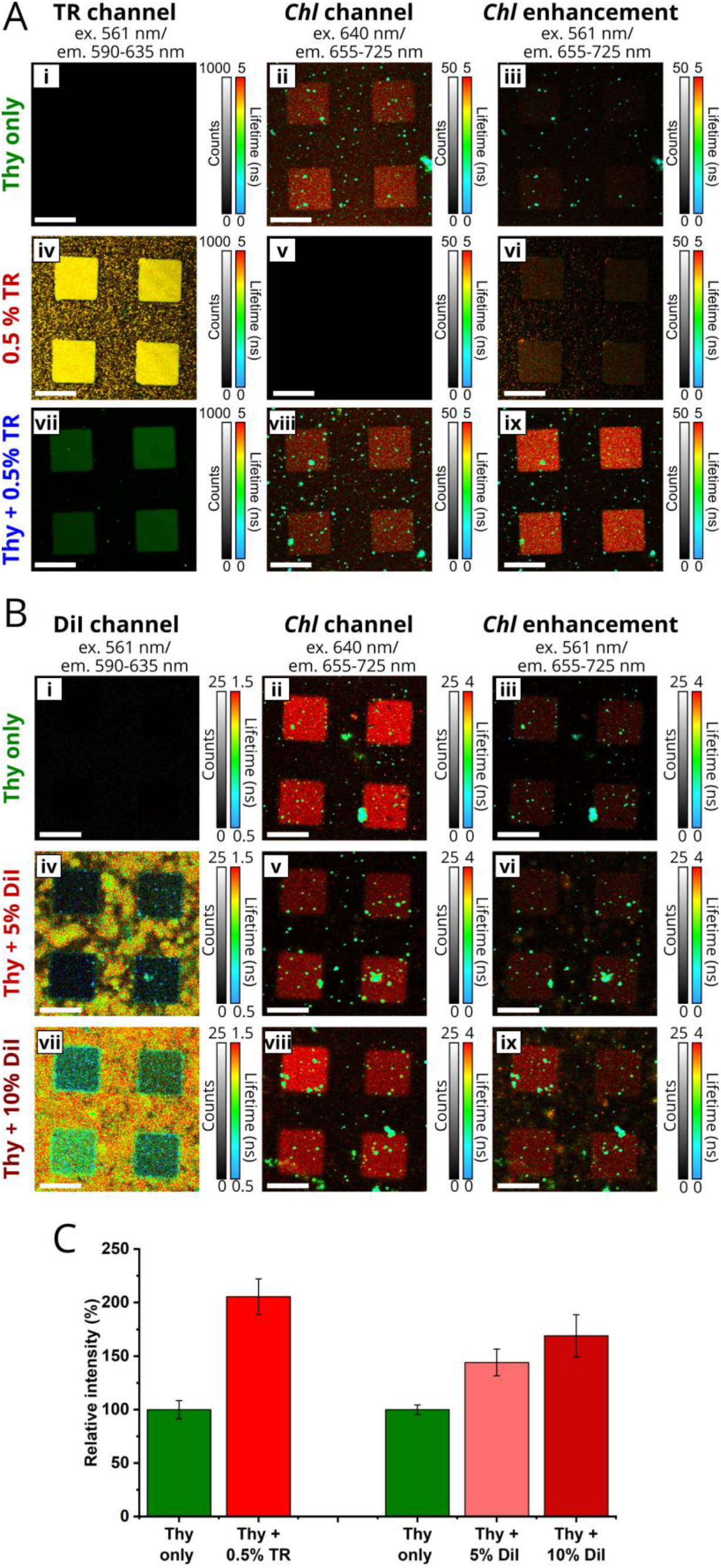
Comparison of the effectiveness of TR and DiI dyes to enhance the fluorescence of surface-supported “hybrid thylakoid membranes”. (**A**) Series of FLIM images to test the effectiveness of TR. “Thy-only” (panels i-iii) represents hybrid membranes generated by fusion of extracted thylakoid membranes and lipid vesicles; “TR-only” (panels iv-vi) represents lipid membranes generated from lipid vesicles containing 0.5% TR-lipids (mol/mol%); “Thy+TR” (panels vii-ix) represents hybrid membranes incorporating TR generated by fusion of extracted thylakoid membranes and lipid vesicles containing 0.5% TR-lipids. Three separate FLIM channels were defined using two alternating lasers and two detectors with selective optical filters to produce the excitation and emission as labelled (ex./em.). Full description of microscopy setup is given in **Methods** (section 2.7). (**B**) Series of FLIM images to test the effectiveness of DiI. The samples are labelled as followed: “Thy-only” (panels i-iii) represents hybrid membranes generated as in (**A**), “Thy+ 5% DiI” (panels iv-vi) and “Thy+ 10% DiI” (panels vii-ix) represent hybrid membranes after the injection of a defined quantity of DiI dissolved in ethanol and incubation for 15 min (0.3 μL or 0.6 μL of DiI at 10 μg/mL) (i.e., DiI injected relative to the estimated lipid on the surface, as mol/mol %). The scale bars in (**A**) and (**B**) represent 20 μm. (**C**) A bar chart comparing the relative intensity in the “Chl enhancement channel” between samples, where the fluorescence in the Thy-only sample is set to 100%. The mean value was calculated from analysis of multiple images from several samples and the error bars represent ± S.D. (see supplementary **Tables S7-S9** for full numerical analysis). Spectral spillover between image channels was minimal but nevertheless carefully accounted for. For all samples, the actual dye concentration generated within membranes cannot be quantified, therefore, the initial amount of dye introduced relative to the known concentration of lipids used is reported (see supplementary **Calculation**). Any excess unincorporated TR-lipids and DiI were removed prior to recording images of the supported membranes, by exchanging the aqueous buffer solution above the membrane multiple times.

Similar FLIM data were acquired to test the effectiveness of DiI for enhancing surface-supported membranes. Here, a hybrid thylakoid membrane sample was assembled onto glass and then DiI was injected into the liquid above this membrane. FLIM images were acquired before and after multiple injections of dye, giving sufficient time between measurements for spontaneous insertion of the DiI into lipid bilayers (**Fig. 6B**). The template pattern was supposed to act as a non-binding surface, however, it did not appear to prevent DiI binding. Images showed that DiI associated non-specifically with both the hybrid membranes (square regions) and the template (meshwork) (**Fig. 6B(iv)** and (**vii**)), however, the DiI located within the hybrid membranes appeared to have a much shorter lifetime (*blue-green* rather than *yellow* colour). This decreased DiI lifetime supports the interpretation of FRET from DiI to LH proteins. As expected, the fluorescence intensity in the standard “Chl channel” was relatively similar before/after DiI injections (**Fig. 6B(ii),(v),(viii)**). A modest increase in the intensity was found in the “Chl enhancement” channel due to the presence of DiI (**Fig. 6A(ix)** compared to (**iii**)).

The extent of the enhancement of Chl fluorescence was calculated for all samples with a careful statistical analysis performed on many corral regions within each sample, see **Fig. 6C**. The emission intensity from thylakoid proteins reached a mean value of 206 ± 17% in the presence of TR, whilst reaching only 169 ± 20% when using the highest concentration of DiI (n = 4, for each), relative to the emission from membranes in the absence of additional dyes. Note that it is difficult to accurately compare the enhancing effect of dyes between hybrid membranes and proteoliposomes because it is challenging to assess the actual dye concentration. Due to the time-consuming nature of FLIM analysis only one concentration of TR was analysed here, but it is known that supported membranes can be formed from liposomes that contain up to 2% (mole/mole) TR-lipids relative to normal lipids [58], so the addition of more TR could allow even greater enhancement of these membranes. However, we may expect a similar lack of membrane stability as observed in proteoliposomes at high dye concentrations. Similarly, a greater quantity of DiI may be injected assuming that the organic solvent used to deliver it does not disrupt the membrane. In summary, lipid-linked dyes appeared to provide superior enhancement effects for surface-associated membranes as compared to injectable dyes, although their full potential remains to be fully investigated.

### 3.7. Benefits and drawbacks: protein-dye self-assembly versus covalent linkages and lipid-linked dyes vs “free” dyes

Our self-assembly approach utilizing lipid membranes as a matrix allowed for *modularity* for choosing different proteins and different dyes and *flexibility* to vary the dye concentration as desired by simply changing the ratio of materials in the starting mixture. Furthermore, compositional modifications were straightforward by the injection of dyes that inserted into pre-existing membranes. Such modularity may be more difficult to generate with alternative approaches employing covalent crosslinking of a dye directly to the LH protein complex [21–24, 61]. Other studies have shown that non-native chromophores may be non-covalently incorporated into the natural pigment binding sites of LH complexes [62–64] by reconstituting the apoprotein with the desired free pigments. However, the new pigments must have suitable chemistry for favourable interactions with binding site residues, making precise control of which pigments can bind challenging [32]. The benefit of using lipid-linked chromophores may be that it relies simply on the consistent interactions between lipid tails (i.e., fatty acyl chains), and only the lipid headgroup is modified to attach the chromophore, allowing a flexible and generally applicable approach to assembly. The drawback is the absence of direct control over the pigment-protein separation distance, leading to a continuum of distances from the lipid-dye to the protein. Consequently, the FRET efficiency will depend on the relative concentrations of each component and on the lateral diffusion of lipids [36, 65]. In summary, both types of coupling strategies can be effective and the choice may depend upon the nanomaterials desired by the researcher.

The dialkylcarbocyanine dyes, DiI and DiR, differed markedly in their effectiveness at enhancing the absorption of LH complexes, presumably related to their ability to insert into lipid membranes or to associate with the protein. DiR suffered from both a low yield of dye incorporation (10-15%) into membranes and a low FRET efficiency (3-11%) of the dye that was present. Therefore, we must assume that structural/chemical issues prevented the occurrence of reasonable coupling between DiR and pigments within LH2. This could be unfavourable lipid-DiR interactions and/or strong DiR-DiR interactions causing DiR aggregation or repulsive interactions (electrostatic or steric) between DiR and LH2. Comparison between of the results between vesicles suspended in solution and membranes deposited onto surfaces also revealed that assembly and insertion effects were critical. DiI appeared to be significantly less effective than TR-lipids at enhancing surface-supported hybrid membranes as compared to a membrane vesicle in solution. Therefore, we can conclude that water-soluble injectable dyes offer greater ease-of-use but can be limited by their ability to insert into lipid bilayers. This necessitates testing the dye of interest for a particular protein and a particular lipid membrane. We would emphasize that even if the FRET efficiency of an injectable dye is lower than a lipid-linked counterpart, their great ease-of-use could make them a useful experimental tool for other researchers. These lipophilic dyes could be injected into any aqueous biological sample of interest and may insert into any membranes present [47]. They could be used to modify natural membranes from photosynthetic organisms and could be useful as either FRET donors or FRET acceptors. They may be applicable to probing whole photosynthetic cells or plant tissues if they can pass cellular barriers to reach their intended destination, which could be achieved with electroporation or similar techniques [66].

## 4. Conclusion

This study provided an in-depth comparison of the relative effectiveness of five different dyes for enhancing the fluorescence of two different LH protein complexes. Our findings show conclusively that the approach of incorporating either lipid-linked dyes or “free” lipophilic dyes into a lipid bilayer together with a membrane protein complex is generally applicable. Lipid-linked Texas Red was able to fill the “green gap” spectral region of minimal natural absorption of plant LHCII, whilst lipid-linked ATTO and Cy7 filled the orange-red and red-NIR spectral ranges of minimal absorption of LH2. By using the highest dye-to-protein ratios, the effective absorption strength of both LH proteins was increased to ~180% of its natural level across the full spectral range or to ~450% when considering the narrower “spectral gap” region. We have also shown that lipophilic dyes, the dialkylcarbocyanines DiI and DiR, will spontaneously insert into pre-formed membranes and enhance an LH complex, although their effectiveness is more variable and dependent on favourable physicochemical interactions. The enhancement of LH complexes was demonstrated both for vesicular membranes in suspension and membranes deposited onto solid surfaces, which may provide interesting opportunities for nanotechnology applications. Indeed, our approach of enhancing bio-mimetic membranes may be compared to concept of dye-sensitized solar cells in which inorganic photoactive surfaces are enhanced with synthetic dyes.

## Acknowledgements

A.M.H. was supported by a studentship from the Engineering and Physical Sciences Research Council (EPSRC, UK) (award number 1807029) and further EPSRC grants (EP/T013958/1 and EP/J017566/1). D.J.K.S. was supported by European Research Council Synergy Award 854126 and Biotechnology and Biological Sciences Research Council (BBSRC, UK) (BB/M000265/1). S.A.M. was supported by a BBSRC studentship (BB/M011151/1). K.M. was supported by the Japan-UK Research Cooperative Program (JPJSBP120195707) and a Grant-in-Aid for Scientific research (Kakenhi) (No. 19H04725 and 21KK0088) from Japan Society for the Promotion of Science (JSPS). C.N.H. was supported by European Research Council Synergy Award 854126. P.G.A. was supported by a University Academic Fellowship from the University of Leeds and a grant from the EPSRC (EP/T013958/1). The PicoQuant FLIM instrument at Leeds was acquired with funding from the BBSRC (BB/R000174/1).

## Reference List

[1] R.E. Blankenship, Molecular mechanisms of photosynthesis, 3rd Ed., Wiley, Hoboken, New Jersey, 2021.

[2] J.D. Rochaix, Regulation and dynamics of the light-harvesting system, Annu. Rev. Plant Biol., 65 (2014) 287–309, 10.1146/annurev-arplant-050213-040226.

[3] H. Lokstein, G. Renger, J.P. Gotze, Photosynthetic light-harvesting (antenna) complexes-structures and functions, Molecules, 26 (2021), 10.3390/molecules26113378.

[4] R. Croce, H. van Amerongen, Light harvesting in oxygenic photosynthesis: Structural biology meets spectroscopy, Science, 369 (2020), 10.1126/science.aay2058.

[5] G.E. Chen, D.P. Canniffe, C.N. Hunter, Three classes of oxygen-dependent cyclase involved in chlorophyll and bacteriochlorophyll biosynthesis, PNAS, 114 (2017) 6280–6285, 10.1073/pnas.1701687114

[6] T.H. Brotosudarmo, A.M. Collins, A. Gall, A.W. Roszak, A.T. Gardiner, R.E. Blankenship, R.J. Cogdell, The light intensity under which cells are grown controls the type of peripheral light-harvesting complexes that are assembled in a purple photosynthetic bacterium, Biochem. J., 440 (2011) 51–61, 10.1042/BJ20110575.

[7] R. Croce, H. van Amerongen, Natural strategies for photosynthetic light harvesting, Nature chemical biology, 10 (2014) 492–501, 10.1038/nchembio.1555.

[8] A. Hitchcock, C.N. Hunter, R. Sobotka, J. Komenda, M. Dann, D. Leister, Redesigning the photosynthetic light reactions to enhance photosynthesis - the PhotoRedesign consortium, Plant J., 109 (2022) 23–34, 10.1111/tpj.15552.

[9] T. Mirkovic, E.E. Ostroumov, J.M. Anna, R. van Grondelle, Govindjee, G.D. Scholes, Light absorption and energy transfer in the antenna complexes of photosynthetic organisms, Chemical reviews, 117 (2017) 249–293, 10.1021/acs.chemrev.6b00002.

[10] V.I. Novoderezhkin, M.A. Palacios, H.V. Amerongen, R. van Grondelle, Energy-transfer dynamics in the LHCII complex of higher plants: modified redfield approach, The journal of physical chemistry. B, 108 (2004) 10363–10375,

[11] M. Sener, J. Strumpfer, J. Hsin, D. Chandler, S. Scheuring, C.N. Hunter, K. Schulten, Forster energy transfer theory as reflected in the structures of photosynthetic light-harvesting systems, Chemphyschem: a European journal of chemical physics and physical chemistry, 12 (2011) 518–531, 10.1002/cphc.201000944.

[12] G. Kodali, J.A. Mancini, L.A. Solomon, T.V. Episova, N. Roach, C.J. Hobbs, P. Wagner, O.A. Mass, K. Aravindu, J.E. Barnsley, K.C. Gordon, D.L. Officer, P.L. Dutton, C.C. Moser, Design and engineering of water-soluble light-harvesting protein maquettes, Chem. Sci., 8 (2017) 316–324, 10.1039/C6SC02417C.

[13] J.A. Mancini, M. Sheehan, G. Kodali, B.Y. Chow, D.A. Bryant, P.L. Dutton, C.C. Moser, De novo synthetic biliprotein design, assembly and excitation energy transfer, J. R. Soc. Interface, 15 (2018), 10.1098/rsif.2018.0021.

[14] W.R. Henson, V.B. Shah, G. Lakin, T. Chadha, H. Liu, R.E. Blankenship, P. Biswas, Production and Performance of a Photosystem I-Based Solar Cell using Nano-Columnar TiO2, Conference paper, (2013), 10.1109/PVSC.2013.6745031.

[15] M. Kamran, J.D. Delgado, V. Friebe, T.J. Aartsma, R.N. Frese, Photosynthetic protein complexes as bio-photovoltaic building blocks retaining a high internal quantum efficiency, Biomacromolecules, 15 (2014) 2833–2838, 10.1021/bm500585s.

[16] V.M. Friebe, J.D. Delgado, D.J.K. Swainsbury, J.M. Gruber, A. Chanaewa, R. van Grondelle, E. von Hauff, D. Millo, M.R. Jones, R.N. Frese, Plasmon-enhanced photocurrent of photosynthetic pigment proteins on nanoporous silver, Adv. Funct. Mater., 26 (2016) 285–292, 10.1002/adfm.201504020.

[17] P.N. Ciesielski, C.J. Faulkner, M.T. Irwin, J.M. Gregory, N.H. Tolk, D.E. Cliffel, G.K. Jennings, Enhanced photocurrent production by Photosystem I multilayer assemblies, Adv. Funct. Mater., 20 (2010) 4048–4054, 10.1002/adfm.201001193.

[18] J. Liu, V.M. Friebe, R.N. Frese, M.R. Jones, Polychromatic solar energy conversion in pigment-protein chimeras that unite the two kingdoms of (bacterio)chlorophyll-based photosynthesis, Nature Communications, 11 (2020) 1542, 10.1038/s41467-020-15321-w.

[19] D.J. Swainsbury, V.M. Friebe, R.N. Frese, M.R. Jones, Evaluation of a biohybrid photoelectrochemical cell employing the purple bacterial reaction centre as a biosensor for herbicides, Biosens. Bioelectron., 58 (2014) 172–178, 10.1016/j.bios.2014.02.050.

[20] A. Ventrella, L. Catucci, T. Placido, F. Longobardi, A. Agostiano, Biomaterials based on photosynthetic membranes as potential sensors for herbicides, Biosensors & Bioelectronics, 26 (2011) 4747–4752, 10.1016/j.bios.2011.05.043.

[21] K. Gundlach, M. Werwie, S. Wiegand, H. Paulsen, Filling the “green gap” of the major lighth-arvesting chlorophyll a/b complex by covalent attachment of Rhodamine Red, Biochimica et biophysica acta, 1787 (2009) 1499–1504, 10.1016/j.bbabio.2009.07.003.

[22] M.A. Harris, J. Jiang, D.M. Niedzwiedzki, J. Jiao, M. Taniguchi, C. Kirmaier, P.A. Loach, D.F. Bocian, J.S. Lindsey, D. Holten, P.S. Parkes-Loach, Versatile design of biohybrid light-harvesting architectures to tune location, density, and spectral coverage of attached synthetic chromophores for enhanced energy capture, Photosynthesis research, 121 (2014) 35–48, 10.1007/s11120-014-9993-8.

[23] Y. Yoneda, T. Noji, T. Katayama, N. Mizutani, D. Komori, M. Nango, H. Miyasaka, S. Itoh, Y. Nagasawa, T. Dewa, Extension of Light-Harvesting Ability of Photosynthetic Light-Harvesting Complex 2 (LH2) through Ultrafast Energy Transfer from Covalently Attached Artificial Chromophores, J. Am. Chem. Soc., 137 (2015) 13121–13129, 10.1021/jacs.5b08508.

[24] J.W. Springer, P.S. Parkes-Loach, K.R. Reddy, M. Krayer, J. Jiao, G.M. Lee, D.M. Niedzwiedzki, M.A. Harris, C. Kirmaier, D.F. Bocian, J.S. Lindsey, D. Holten, P.A. Loach, Biohybrid photosynthetic antenna complexes for enhanced light-harvesting, J. Am. Chem. Soc., 134 (2012) 4589–4599, 10.1021/ja207390y.

[25] W. Li, S. Wu, H. Zhang, X. Zhang, J. Zhuang, C. Hu, Y. Liu, B. Lei, L. Ma, X. Wang, Enhanced biological photosynthetic efficiency using light-harvesting engineering with dual-emissive carbon dots, Adv. Funct. Mater., 28 (2018) 1804004, 10.1002/adfm.201804004.

[26] G. Amoruso, J. Liu, D.W. Polak, K. Tiwari, M.R. Jones, T.A.A. Oliver, High-efficiency excitation energy transfer in biohybrid quantum dot-bacterial reaction center nanoconjugates, J. Phys. Chem. Lett., 12 (2021) 5448–5455, 10.1021/acs.jpclett.1c01407.

[27] F.J. Schmitt, E.G. Maksimov, P. Hätti, J. Weißenborn, V. Jeyasangar, A.P. Razjivin, V.Z. Paschenko, T. Friedrich, G. Renger, Coupling of different isolated photosynthetic light harvesting complexes and CdSe/ZnS nanocrystals via Förster resonance energy transfer, Biochimica et biophysica acta, 1817 (2012) 1461–1470, 10.1016/j.bbabio.2012.03.030.

[28] K.J. Grayson, K.M. Faries, X. Huang, P. Qian, P. Dilbeck, E.C. Martin, A. Hitchcock, C. Vasilev, J.M. Yuen, D.M. Niedzwiedzki, G.J. Leggett, D. Holten, C. Kirmaier, C. Neil Hunter, Augmenting light coverage for photosynthesis through YFP-enhanced charge separation at the Rhodobacter sphaeroides reaction centre, Nat. Commun., 8 (2017) 13972, 10.1038/ncomms13972.

[29] G. Amoruso, J. Liu, D.W. Polak, K. Tiwari, M.R. Jones, T.A.A. Oliver, High-Efficiency Excitation Energy Transfer in Biohybrid Quantum Dot-Bacterial Reaction Center Nanoconjugates, The Journal of Physical Chemistry Letters, 12 (2021) 5448–5455, 10.1021/acs.jpclett.1c01407.

[30] P.G. Adams, C. Vasilev, C.N. Hunter, M.P. Johnson, Correlated fluorescence quenching and topographic mapping of Light-Harvesting Complex II within surface-assembled aggregates and lipid bilayers, Biochimica et biophysica acta, 1859 (2018) 1075–1085, 10.1016/j.bbabio.2018.06.011.

[31] P. Qian, D.J.K. Swainsbury, T.I. Croll, P. Castro-Hartmann, G. Divitini, K. Sader, C.N. Hunter, Cryo-EM structure of the *Rhodobacter sphaeroides* Light-Harvesting 2 Complex at 2.1 A, Biochemistry, 60 (2021) 3302–3314, 10.1021/acs.biochem.1c00576.

[32] D.J.K. Swainsbury, K.M. Faries, D.M. Niedzwiedzki, E.C. Martin, A.J. Flinders, D.P. Canniffe, G. Shen, D.A. Bryant, C. Kirmaier, D. Holten, C.N. Hunter, Engineering of B800 bacteriochlorophyll binding site specificity in the *Rhodobacter sphaeroides* LH2 antenna, Biochimica et biophysica acta, 1860 (2019) 209–223, 10.1016/j.bbabio.2018.11.008.

[33] M. Son, A. Pinnola, S.C. Gordon, R. Bassi, G.S. Schlau-Cohen, Observation of dissipative chlorophyll-to-carotenoid energy transfer in light-harvesting complex II in membrane nanodiscs, Nat. Commun., 11 (2020) 1295, 10.1038/s41467-020-15074-6.

[34] A. Pandit, N. Shirzad-Wasei, L.M. Wlodarczyk, H. van Roon, E.J. Boekema, J.P. Dekker, W.J. de Grip, Assembly of the major light-harvesting complex II in lipid nanodiscs, Biophys. J., 101 (2011) 2507–2515, 10.1016/j.bpj.2011.09.055.

[35] Y. Yoneda, M. Kito, D. Mori, A. Goto, M. Kondo, H. Miyasaka, Y. Nagasawa, T. Dewa, Ultrafast energy transfer between self-assembled fluorophore and photosynthetic light-harvesting complex 2 (LH2) in lipid bilayer, The Journal of Chemical Physics, 156 (2022) 095101, 10.1063/5.0077910.

[36] A.M. Hancock, S.A. Meredith, S.D. Connell, L.J.C. Jeuken, P.G. Adams, Proteoliposomes as energy transferring nanomaterials: enhancing the spectral range of light-harvesting proteins using lipid-linked chromophores, Nanoscale, 11 (2019) 16284–16292, 10.1039/C9NR04653D.

[37] P. Yuan, D. Walt, Calculation for Fluorescence Modulation by Absorbing Species and Its Application to Measurements Using Optical Fibers, Anal. Chem., 59 (1957) 2391–2394,

[38] T. Yoneda, Y. Tanimoto, D. Takagi, K. Morigaki, Photosynthetic model membranes of natural plant thylakoid embedded in a patterned polymeric lipid bilayer, Langmuir, 36 (2020) 5863–5871, 10.1021/acs.langmuir.0c00613.

[39] S.A. Meredith, T. Yoneda, A.M. Hancock, S.D. Connell, S.D. Evans, K. Morigaki, P.G. Adams, Model lipid membranes assembled from natural plant thylakoids into 2D microarray patterns as a platform to assess the organization and photophysics of light-harvesting proteins, Small, 17 (2021) e2006608, 10.1002/smll.202006608.

[40] Z. Liu, H. Yan, K. Wang, T. Kuang, J. Zhang, L. Gui, X. An, W. Chang, Crystal structure of spinach major light-harvesting complex at 2.72 A resolution, Nature, 428 (2004) 287–292, 10.1038/nature02373.

[41] J. Standfuss, A.C.T. van Scheltinga, M. Lamborghini, W. Kuhlbrandt, Mechanisms of photoprotection and nonphotochemical quenching in pea light-harvesting complex at 2.5A resolution, Embo Journal, 24 (2005) 919–928, 10.1038/sj.emboj.7600585.

[42] A.M. Hancock, M. Son, M. Nairat, T. Wei, L.J.C. Jeuken, C.D.P. Duffy, G.S. Schlau-Cohen, P.G. Adams, Ultrafast energy transfer between lipid-linked chromophores and plant light-harvesting complex II, Chemphyschem: a European journal of chemical physics and physical chemistry, 23 (2021) 19511–19524, 10.1039/d1cp01628h.

[43] D.M. Niedzwiedzki, D.J.K. Swainsbury, D.P. Canniffe, C.N. Hunter, A. Hitchcock, A photosynthetic antenna complex foregoes unity carotenoid-to-bacteriochlorophyll energy transfer efficiency to ensure photoprotection, PNAS, 117 (2020) 6502–6508, 10.1073/pnas.1920923117.

[44] S. Hess, K.J. Visscher, T. Pullerits, V. Sundstroem, G.J.S. Fowler, C.N. Hunter, Enhanced rates of subpicosecond energy transfer in blue-shifted light-harvesting LH2 mutants of Rhodobacter sphaeroides, Biochemistry, 33 (1994) 8300–8305, 10.1021/bi00193a017.

[45] G.J.S. Fowler, R.W. Visschers, G.G. Grief, R. van Grondelle, C.N. Hunter, Genetically modified photosynthetic antenna complexes with blueshifted absorbance bands, Nature, 355 (1992) 848–850, 10.1038/355848a0.

[46] P. Qian, T.I. Croll, A. Hitchcock, P.J. Jackson, J.H. Salisbury, P. Castro-Hartmann, K. Sader, D.J.K. Swainsbury, C.N. Hunter, Cryo-EM structure of the dimeric Rhodobacter sphaeroides RC-LH1 core complex at 2.9 A: the structural basis for dimerisation, Biophys. J., 478 (2021) 3923–3937, 10.1042/BCJ20210696.

[47] R.D. Klausner, E.D. Wolf, Selectivity of fluorescent lipid analogs for lipid domains, Biochemistry, 19 (1980) 6199–6203,

[48] P.G. Adams, D.J. Mothersole, I.W. Ng, J.D. Olsen, C.N. Hunter, Monomeric RC-LH1 core complexes retard LH2 assembly and intracytoplasmic membrane formation in PufX-minus mutants of *Rhodobacter sphaeroides*, BBA Bioenerg., 1807 (2011) 1044–1055, 10.1016/j.bbabio.2011.05.019.

[49] S. Scheuring, J.N. Sturgis, Atomic force microscopy of the bacterial photosynthetic apparatus: plain pictures of an elaborate machinery, Photosynth Res, 102 (2009) 197–211, 10.1007/s11120-009-9413-7.

[50] S. Bahatyrova, R.N. Frese, K.O. van der Werf, C. Otto, C.N. Hunter, J.D. Olsen, Flexibility and size heterogeneity of the LH1 light harvesting complex revealed by atomic force microscopy: functional significance for bacterial photosynthesis, J. Biol. Chem., 279 (2004) 21327–21333, 10.1074/jbc.M313039200.

[51] M. Bally, K. Bailey, K. Sugihara, D. Grieshaber, J. Voros, B. Stadler, Liposome and lipid bilayer arrays towards biosensing applications, Small, 6 (2010) 2481–2497, 10.1002/smll.201000644.

[52] T. Noji, M. Matsuo, N. Takeda, A. Sumino, M. Kondo, M. Nango, S. Itoh, T. Dewa, Lipid-Controlled Stabilization of Charge-Separated States (P+QB-) and Photocurrent Generation Activity of a Light-Harvesting-Reaction Center Core Complex (LH1-RC) from Rhodopseudomonas palustris, The Journal of Physical Chemistry B, 122 (2018) 1066–1080, 10.1021/acs.jpcb.7b09973.

[53] T. Förster, Delocalized excitation and excitation transfer, Modern Quantum Chemistry Istanbul Lectures, 3 (1965) 93–137,

[54] I. AAT Bioquest, Quest Database™ Extinction Coefficient [Texas Red]. Accessed 12 Jul. 2022, in, https://www.aatbio.com/resources/extinction-coefficient/texas_red_sulforhodamine_101_sulfonyl_chloride, 2022,

[55] J. Gravier, F.P. Navarro, T. Delmas, F. Mittler, A.C. Couffin, F. Vinet, I. Texier, Lipidots: competitive organic alternative to quantum dots for in vivo fluorescence imaging, J. Biomed. Opt., 16 (2011) 096013, 10.1117/1.3625405.

[56] ATTO-TEC, Product Information: ATTO 647N, in: Accessed 12 Jul. 2022, https://www.atto-tec.com/fileadmin/user_upload/Katalog_Flyer_Support/ATTO_647N.pdf, 2019,

[57] I. AAT Bioquest, Quest Database™ Extinction Coefficient [Cy7 (Cyanine-7)]. Accessed 12 Jul. 2022, in, https://www.aatbio.com/resources/extinction-coefficient/cy_7cyanine7, 2022,

[58] J.A. Titus, R. Haugland, S.O. Sharrow, D.M. Segal, Texas red, a hydrophilic, red-emitting flourophore for use with flourescein in dual parameter flow microfluorometric and fluorescence microscopic studies, Journal of Immunological Methods, 50 (1982) 193–204, 10.1016/0022-1759(82)90225-3.

[59] Y. Yoneda, D. Kato, M. Kondo, K.V.P. Nagashima, H. Miyasaka, Y. Nagasawa, T. Dewa, Sequential energy transfer driven by monoexponential dynamics in a biohybrid light-harvesting complex 2 (LH2), Photosynth Res, 143 (2020) 115–128, 10.1007/s11120-019-00677-y.

[60] Y. Yoneda, T. Noji, T. Katayama, N. Mizutani, D. Komori, M. Nango, H. Miyasaka, S. Itoh, Y. Nagasawa, T. Dewa, Extension of Light-Harvesting Ability of Photosynthetic Light-Harvesting Complex 2 (LH2) through Ultrafast Energy Transfer from Covalently Attached Artificial Chromophores, J Am Chem Soc, 137 (2015) 13121–13129, 10.1021/jacs.5b08508.

[61] P.K. Dutta, S. Lin, A. Loskutov, S. Levenberg, D. Jun, R. Saer, J.T. Beatty, Y. Liu, H. Yan, N.W. Woodbury, Reengineering the optical absorption cross-section of photosynthetic reaction centers, J. Am. Chem. Soc., 136 (2014) 4599–4604, 10.1021/ja411843k.

[62] E. Elias, N. Liguori, Y. Saga, J. Schafers, R. Croce, Harvesting far-red light with plant antenna complexes incorporating Chlorophyll *d*, Biomacromolecules, 22 (2021) 3313–3322, 10.1021/acs.biomac.1c00435.

[63] J.L. Herek, N.J. Fraser, T. Pullerits, P. Martinsson, T. Polívka, H. Scheer, R.J. Cogdell, V. Sundström, B800→B850 Energy transfer mechanism in bacterial LH2 complexes investigated by B800 pigment exchange, Biophys. J., 78 (2000) 2590–2596, 10.1016/S0006-3495(00)76803-2.

[64] S.C. Chi, D.J. Mothersole, P. Dilbeck, D.M. Niedzwiedzki, H. Zhang, P. Qian, C. Vasilev, K.J. Grayson, P.J. Jackson, E.C. Martin, Y. Li, D. Holten, C. Neil Hunter, Assembly of functional photosystem complexes in *Rhodobacter sphaeroides* incorporating carotenoids from the spirilloxanthin pathway, Biochimica et biophysica acta, 1847 (2015) 189–201, 10.1016/j.bbabio.2014.10.004.

[65] A.M. Hancock, M. Son, M. Nairat, T. Wei, L.J.C. Jeuken, C.D.P. Duffy, G.S. Schlau-Cohen, P.G. Adams, Ultrafast energy transfer between lipid-linked chromophores and plant light-harvesting complex II, Physical Chemistry Chemical Physics, 23 (2021) 19511–19524, 10.1039/D1CP01628H.

[66] Y. Rosemberg, R. Korenstein, Electroporation of the photosynthetic membrane: A study by intrinsic and external optical probes., Biophys. J., 58 (1990) 823–832, 10.1016/S0006-3495(90)82428-0.

