## Supplementary Information for "Enhancing the spectral range of plant and bacterial Light-Harvesting pigment-protein complexes with various synthetic chromophores incorporated into lipid vesicles"

### Electronic supplementary information (ESI)

**ESI 1 – Gel electrophoresis of LH pigment-protein complexes**

Trimeric LHCII was purified as described in main-text **Methods** section 2.2 and purity was confirmed by polyacrylamide gel electrophoresis (PAGE), shown in **Figure S1** below. Single strong bands in both SDS and native PAGE gels for post-SEC (post-size exclusion chromatography) LHCII trimers demonstrate protein purity [1]. LHCII trimer molar concentration is estimated as [Chl mM concentration] x 42, where "chlorophyll mM concentration" is determined from absorbance measurements after methanol/acetone pigment extraction as in [2]. This leads to a calculated molar extinction coefficient value of 1,846,040 $M^{-1}cm^{-1}$ at 675 nm allowing conversion from absorbance to molar concentration. The Chl a/b ratio from methanol/acetone pigment extraction of purified trimeric LHCII was typically 1.4.

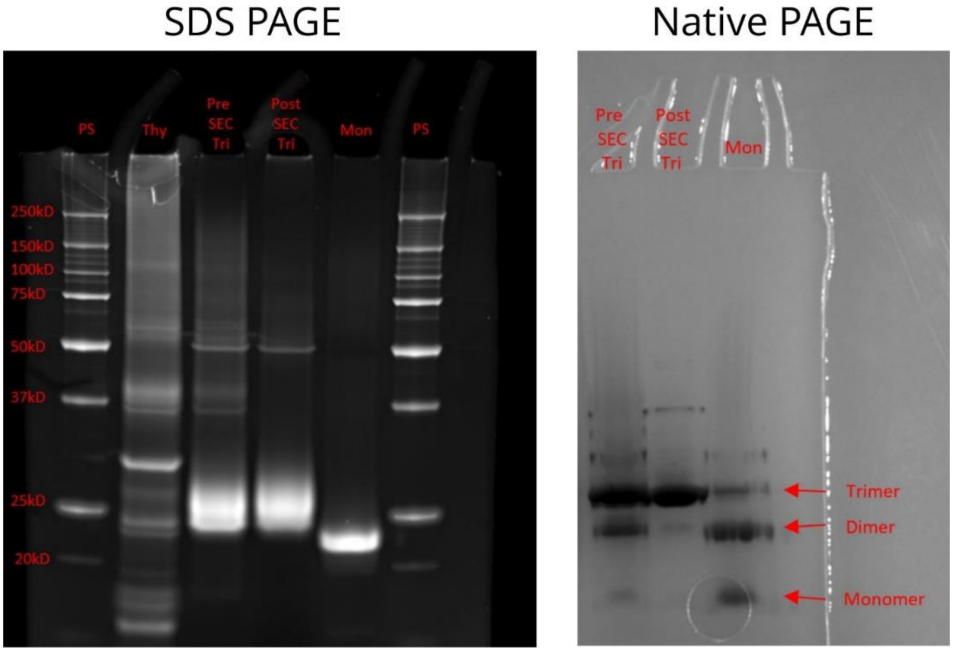


**Fig. S1** *Left panel: SDS-PAGE gel with SYPRO-RUBY stain. Gel lanes show, in order: protein standard (PS), Extracted thylakoid membrane, i.e., detergent solubilised thylakoids before being sucrose gradient purification (Thy), LHCII "before FPLC" containing impurities, i.e., before size exclusion chromatography (pre SEC Tri), purified trimeric LHCII i.e., trimer fractions after size exclusion chromatography (Post SEC Tri), purified monomeric LHCII, i.e., monomer fractions after size exclusion chromatography (Mon). Right panel: Native-PAGE gel (right panel) was ran at 4°C and then stained with Coomassie. Native gel lanes show, in order: LHCII "before FPLC", purified trimeric LHCII, and purified monomeric LHCII, as above.*

#### ESI 2 – Theory: Full expressions for photophysical properties and calculations of Forster radius (*R₀*) and spectral overlap integral (*J*)

Absorbance is quantified as a function of wavelength ($\lambda$) as:

$$A_{dye}(\lambda) = \varepsilon(\lambda) \cdot C \cdot l \qquad \textit{(Eqn. S1)}$$

where, $\varepsilon(\lambda)$ is the molar absorption coefficient at a given wavelength (a measure of the absorbing strength of a material), $C$ is the molar concentration and $l$ is the pathlength through which the light beam passes (i.e., the thickness of the cuvette).

Assuming that the donor is successfully promoted to an excited state, the FRET efficiency ($E_{FRET}$), defined as the quantum yield of energy transfer process over all other possible decay processes (fluorescence and non-radiative decay). This depends upon the donor-acceptor pair having a strong dipole-dipole coupling and also a good energetic overlap which depends strongly upon the donor-acceptor separation distance ($r$) and their inherent photophysical properties. $E_{FRET}$ can then be related to $r$ by applying a simple form of Förster theory [3]:

$$E_{FRET} = \frac{1}{1+\left(\frac{r}{R_0}\right)^6} \qquad \textit{(Eqn. S2)}$$

where, $R_0$ is known as the Förster radius for the donor-acceptor pair. This is defined as the distance where energy transfer is 50 % efficient [3] and can be calculated as:

$$R_0^6 = 8.79 \times 10^{-5} J\, \kappa^2\, n^4\, \phi_D \qquad \textit{(Eqn. S3)}$$

where $R_0$ is the Förster distance in Å; *J* is the spectral overlap integral; $\kappa$ is the orientation factor; *n* is the refractive index of optical medium; $\phi_D$ is the fluorescence quantum yield of the donor.

The *J* factor can be calculated from the overlap of the normalised donor emission spectrum and acceptor absorption spectra using the following equation:

$$J = \int_0^{\infty} (F_D(\lambda)\, \varepsilon_A(\lambda)\, \lambda^4)\, d\lambda \qquad \textit{(Eqn. S4)}$$

where $F_D(\lambda)$ is the normalized fluorescence emission spectrum of the donor, $\varepsilon_A(\lambda)$ is the molar absorption coefficient of the acceptor as a function of wavelength (obtained from an absorption spectrum of the acceptor) and $\lambda$ is the wavelength. Whilst the protein is not a simple acceptor molecule, as excitation energy may be passed from the donor dye to any of the carotenoid or (B)Chl pigments within the protein it is reasonable as a rough approximation to use the protein's absorption spectrum to represent the sum of the many pigments within the LH protein complex.

Values for *J* and $R_0$ listed in the main text **Table 1** were calculated for each dye-protein combination as follows. *J* was calculated using *Eqn. S4* by inputting the appropriate donor emission spectrum and acceptor absorption spectra as shown in **Fig. S2A**. $R_0$ was then calculated using *Eqn. S3*, using these values for this *J*. In the current study, we assumed that the relative orientation of the transition dipoles of donor and acceptor molecules was random, equating to $\kappa^2$ = 2/3 [4]. The refractive index (*n*) of the optical medium was assumed to be halfway between that of water (1.33) and lipid tail groups (1.55) [5] with a value of 1.45. The resulting *J* integrals for each dye-protein combination are shown in **Fig. S2B**.

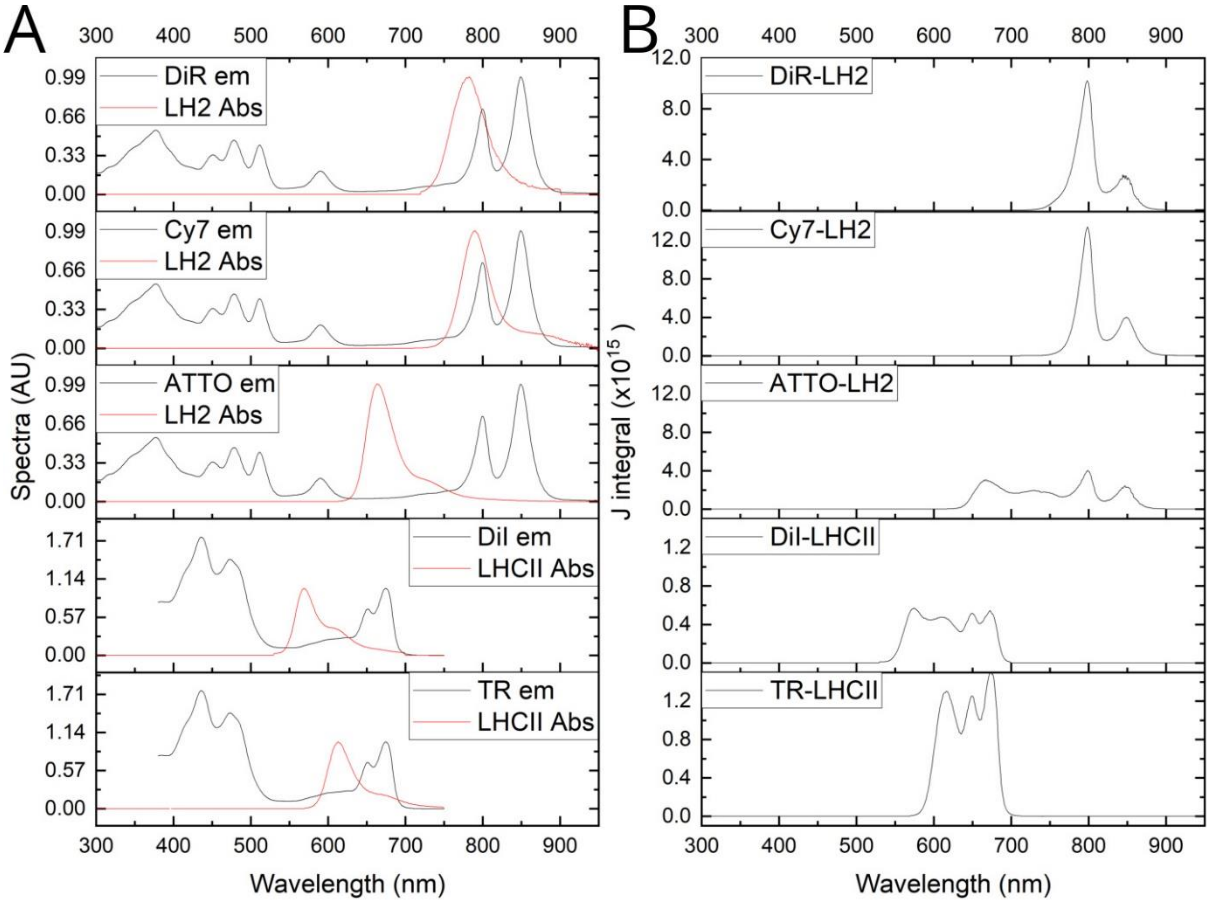


**Fig. S2.** *(A) Normalised donor emission and acceptor emission spectra for each dye-protein combination. (B) Calculated integrated spectral overlap (J integral) for each dye-protein combination.*

**ESI 3 – Analysis of absorbance spectra and calculation of concentrations for each dye-protein combination**

The concentration of protein/dye in samples was determined by measuring the sample absorbance and applying the molar absorption coefficient of each component, as described below. For plant LHCII and bacterial LH2 the molar absorption coefficients are 1,846,040 $M^{-1}cm^{-1}$ (at 675 nm) and 2,999,990 $M^{-1}cm^{-1}$ (at 850 nm), respectively (as described in the main text **Methods** section 2.5). Note that this relates to the trimeric form of plant LHCII. Molar absorption coefficients for each dye are displayed in main text **Table 1**.

The values measured for dye absorbance were corrected for their overlap with the absorbance from the LH protein by subtracting the known fractional overlap at wavelengths used for calculating the concentration. Relative to its absorption peak at 673-675 nm plant LHCII has a contribution of 24.3% at ~582.5 nm and 15.2% at ~555 nm; this is taken into account for calculating TR and DiI absorption, respectively. Relative to its absorption peak at ~850 nm bacterial LH2 has a contribution of 13.3% at ~765 nm, 1.1% at ~646 nm, and 6.4% at ~750 nm; this is taken into account for calculating Cy7, ATTO, and DiR concentrations, respectively.

The concentrations calculated of LH proteins and dyes within proteoliposome samples are shown below in **Table S1**. The concentration of lipids was 1 mM in all proteoliposomes.

| Sample Name (initial conc.) | LHCII Abs | LHCII Conc. μM | TR Abs | TR Abs corr. | TR conc. μM | LHCII Yield % | TR Yield % | Dye:LHCII Ratio | Dye:Chl Ratio |
|---|---|---|---|---|---|---|---|---|---|
| 0.7μM LHCII* | 1.24 | 0.67 | 0.30 | 0.00 | 0.00 | 96 | N/A | 0.0 | 0.00 |
| 0.7μM LHCII + 15μM TR* | 1.15 | 0.62 | 1.27 | 0.99 | 11.71 | 89 | 78 | 13.7 | 0.33 |
| 0.7μM LHCII + 22.5μM TR** | 1.06 | 0.57 | 1.90 | 1.64 | 19.38 | 82 | 86 | 24.6 | 0.59 |
| 0.7μM LHCII + 30μM TR** | 1.12 | 0.61 | 2.42 | 2.15 | 25.38 | 87 | 85 | 30.5 | 0.73 |
| 0.7μM LHCII + 37.5μM TR** | 1.13 | 0.61 | 2.88 | 2.61 | 30.83 | 87 | 82 | 36.8 | 0.88 |
| 0.7μM LHCII + 75μM TR*** | 1.10 | 0.59 | 5.54 | 5.27 | 62.33 | 85 | 83 | 76.5 | 1.82 |

| Sample Name (initial conc.) | LHCII Abs | LHCII Conc. μM | DiI Abs | DiI Abs Corr. | DiI Conc. μM | LHCII Yield % | DiI Yield % | Dye:LHCII Ratio | Dye:Chl Ratio |
|---|---|---|---|---|---|---|---|---|---|
| 0.7μM LHCII* | 0.75 | 0.41 | 0.11 | 0.00 | 0.00 | 58 | N/A | 0.0 | 0.00 |
| 0.7μM LHCII + 8μM* | 0.70 | 0.38 | 0.44 | 0.39 | 2.69 | 54 | 34 | 14.8 | 0.35 |
| 0.7μM LHCII + 12μM DiI* | 0.69 | 0.37 | 0.86 | 0.80 | 5.54 | 53 | 46 | 21.8 | 0.52 |
| 0.7μM LHCII + 20μM DiI* | 0.66 | 0.36 | 1.18 | 1.12 | 7.79 | 51 | 39 | 38.4 | 0.91 |
| 0.7μM LHCII + 10μM DiI* | 0.60 | 0.33 | 1.85 | 1.80 | 12.49 | 46 | 125 | 13.5 | 0.32 |
| 0.7μM LHCII + 20μM DiI** | 0.61 | 0.33 | 1.50 | 1.45 | 10.07 | 47 | 50 | 30.7 | 0.73 |
| 0.7μM LHCII + 30μM DiI** | 0.57 | 0.31 | 2.06 | 2.01 | 13.97 | 44 | 47 | 45.2 | 1.08 |
| 0.7μM LHCII + 40μM DiI** | 0.55 | 0.30 | 2.83 | 2.79 | 19.35 | 42 | 48 | 65.4 | 1.56 |
| 0.7μM LHCII + 50μM DiI** | 0.51 | 0.28 | 3.65 | 3.61 | 25.04 | 39 | 50 | 90.6 | 2.16 |
| 0.7μM LHCII + 75μM DiI*** | 0.61 | 0.33 | 6.50 | 6.45 | 44.82 | 47 | 60 | 135.2 | 3.22 |
| 0.7μM LHCII + 100μM DiI*** | 0.59 | 0.32 | 8.48 | 8.43 | 58.54 | 46 | 59 | 183.8 | 4.38 |

| Sample<br>Name (initial conc.) | LH2 Abs | LH2 Conc.<br>µM | ATTO Abs | ATTO Abs Corr. | ATTO Conc.<br>µM | LH2 Yield<br>% | ATTO Yield<br>% | Dye:LH2 Ratio | Dye:BChl Ratio |
|---|---|---|---|---|---|---|---|---|---|
| 0.5µM LH2* | 1.26 | 0.42 | 0.01 | 0.00 | 0.0 | 84 | N/A | 0.0 | 0.00 |
| 0.5µM LH2 + 2µM ATTO* | 1.58 | 0.53 | 0.23 | 0.21 | 1.4 | 106 | 69 | 2.6 | 0.10 |
| 0.5µM LH2 + 4µM ATTO* | 1.49 | 0.50 | 0.48 | 0.46 | 3.1 | 100 | 77 | 6.2 | 0.23 |
| 0.5µM LH2 + 6µM ATTO* | 1.36 | 0.45 | 0.91 | 0.90 | 6.0 | 91 | 100 | 13.2 | 0.49 |
| 0.5µM LH2 + 8µM ATTO* | 1.44 | 0.48 | 0.75 | 0.73 | 4.9 | 96 | 61 | 10.2 | 0.38 |
| 0.5µM LH2 + 10µM ATTO* | 1.40 | 0.47 | 1.26 | 1.25 | 8.3 | 94 | 83 | 17.8 | 0.66 |
| 0.5µM LH2 + 12µM ATTO* | 1.38 | 0.46 | 1.31 | 1.30 | 8.6 | 92 | 72 | 18.8 | 0.70 |

| Sample<br>Name (initial conc.) | LH2 Abs | LH2 Conc.<br>µM | Cy7 Abs | Cy7 Abs Corr. | Cy7 Conc.<br>µM | LH2 Yield<br>% | Cy7 Yield<br>% | Dye:LH2 Ratio | Dye:BChl Ratio |
|---|---|---|---|---|---|---|---|---|---|
| 0.5µM LH2* | 1.72 | 0.52 | 0.22 | -0.01 | 0.00 | 103 | N/A | 0.0 | 0.00 |
| 0.5µM LH2 + 1.5µM Cy7* | 1.46 | 0.44 | 0.46 | 0.27 | 1.08 | 88 | 68 | 2.2 | 0.08 |
| 0.5µM LH2 + 3.0µM Cy7* | 1.32 | 0.39 | 0.68 | 0.51 | 2.03 | 79 | 63 | 4.6 | 0.17 |
| 0.5µM LH2 + 4.5µM Cy7* | 1.38 | 0.41 | 0.89 | 0.71 | 2.84 | 83 | 78 | 6.2 | 0.23 |
| 0.5µM LH2 + 6.0µM Cy7* | 1.67 | 0.50 | 1.39 | 1.17 | 4.68 | 100 | 73 | 8.4 | 0.31 |
| 0.5µM LH2 + 7.5µM Cy7* | 1.65 | 0.49 | 1.59 | 1.37 | 5.49 | 99 | 69 | 9.9 | 0.37 |
| 0.5µM LH2 + 9.0µM Cy7* | 1.45 | 0.43 | 1.75 | 1.56 | 6.25 | 87 | 69 | 12.9 | 0.48 |

| Sample Name (initial conc.) | LH2 Abs | LH2 Conc.<br>µM | ATTO Abs | ATTO Abs Corr. | ATTO Conc.<br>µM | Cy7 Abs | Cy7 Abs Corr. | Cy7 Conc.<br>µM | LH2 Yield<br>% | ATTO Yield<br>% | Cy7 Yield<br>% | Dye: LH2 Ratio | Dye: BChl Ratio |
|---|---|---|---|---|---|---|---|---|---|---|---|---|---|
| 0.5µM LH2* | 1.32 | 0.44 | 0.03 | 0.00 | 0.01 | 0.13 | 0.00 | 0.00 | 88 | N/A | N/A | 0.0 | 0.00 |
| 0.5µM LH2 + 6µM ATTO + 4.5µM Cy7* | 1.33 | 0.44 | 0.57 | 0.44 | 2.93 | 0.62 | 0.48 | 1.93 | 89 | 49 | 43 | 11.0 | 0.41 |
| 0.5µM LH2 + 6µM ATTO + 9µM Cy7* | 1.34 | 0.45 | 0.65 | 0.37 | 2.44 | 1.34 | 1.21 | 4.83 | 89 | 41 | 54 | 16.3 | 0.60 |
| 0.5µM LH2 + 12µM ATTO + 4.5µM Cy7* | 1.32 | 0.44 | 1.19 | 1.04 | 6.95 | 0.71 | 0.58 | 2.30 | 88 | 58 | 51 | 21.1 | 0.78 |
| 0.5µM LH2 + 12µM ATTO + 9µM Cy7* | 1.32 | 0.44 | 1.27 | 0.97 | 6.47 | 1.39 | 1.26 | 5.03 | 88 | 54 | 56 | 26.1 | 0.97 |

| Sample<br>Name (initial conc.) | LH2 Abs | LH2 Conc.<br>µM | DiR Abs | DiR Abs Corr. | DiR Conc.<br>µM | LH2 Yield<br>% | DiR Yield<br>% | Dye:LH2 Ratio | Dye:BChl Ratio |
|---|---|---|---|---|---|---|---|---|---|
| 0.5µM LH2 + 0µM DiR* | 1.27 | 0.42 | 0.08 | 0.00 | 0.0 | 85 | N/A | 0.0 | 0.00 |
| 0.5µM LH2 + 4µM DiR* | 1.31 | 0.44 | 0.24 | 0.16 | 0.6 | 88 | 14 | 1.3 | 0.05 |
| 0.5µM LH2 + 8µM DiR* | 1.32 | 0.44 | 0.33 | 0.24 | 0.9 | 88 | 11 | 2.0 | 0.07 |
| 0.5µM LH2 + 12µM DiR* | 1.32 | 0.44 | 0.41 | 0.32 | 1.2 | 88 | 10 | 2.7 | 0.10 |
| 0.5µM LH2 + 16µM DiR* | 1.32 | 0.44 | 0.55 | 0.46 | 1.7 | 88 | 11 | 3.9 | 0.15 |
| 0.5µM LH2 + 20µM DiR* | 1.31 | 0.44 | 0.65 | 0.57 | 2.1 | 87 | 11 | 4.8 | 0.18 |
| 0.5µM LH2 + 30µM DiR* | 1.54 | 0.51 | 0.97 | 0.87 | 3.2 | 102 | 11 | 6.3 | 0.23 |
| 0.5µM LH2 + 40µM DiR* | 1.55 | 0.52 | 1.19 | 1.09 | 4.0 | 104 | 10 | 7.8 | 0.29 |
| 0.5µM LH2 + 50µM DiR* | 1.56 | 0.52 | 1.55 | 1.45 | 5.4 | 104 | 11 | 10.3 | 0.38 |
| 0.5µM LH2 + 60µM DiR* | 1.57 | 0.52 | 1.86 | 1.76 | 6.5 | 105 | 11 | 12.4 | 0.46 |
| 0.5µM LH2 + 70µM DiR* | 1.57 | 0.52 | 2.17 | 2.07 | 7.6 | 105 | 11 | 14.6 | 0.54 |
| 0.5µM LH2 + 80µM DiR* | 1.60 | 0.53 | 2.41 | 2.31 | 8.5 | 107 | 11 | 16.0 | 0.59 |
| 0.5µM LH2 + 90µM DiR* | 1.61 | 0.54 | 2.73 | 2.63 | 9.7 | 108 | 11 | 18.1 | 0.67 |
| 0.5µM LH2 + 100µM DiR* | 1.62 | 0.54 | 3.07 | 2.97 | 11.0 | 108 | 11 | 20.4 | 0.75 |

**Table S1** *Absorption and calculated concentrations of LH proteins and dyes for all samples.*

*Abs = Absorbance at extinction coefficient peak position;*

*Conc. = molar concentration. The molar concentrations for each dye and protein were calculated using the molar absorption coefficients as noted in the text above.*

*Abs Corr. = Absorbance corrected for spectral overlap.*

*Yield was defined as the calculated concentration divided by the initial concentration in the starting mixture.*

*Samples were diluted before cuvette-based spectroscopy by the dilution factors indicated: * = x15, ** = x30, *** = x60*

**ESI 4 - Analysis of Texas Red only DOPC liposomes showing self-quenching**

Self-quenching of TR-DHPE in liposomes at higher concentrations is a known effect that has been established in previous publications [6]. In order to assess this effect, liposomes containing 1 mM lipid were prepared with varying concentrations of TR (**Fig. S3A**). To determine the level of fluorescence self-quenching we compared the relative fluorescence emission-to-absorbance ratio of the samples. This intensity value allows one to compare the level of emission per mole of TR at different concentrations (**Fig. S3B** and **Table S2**). As the TR concentration in liposomes increases from 4.7µM to 7.7µM there is a 20% decrease in the relative fluorescence emission confirming that TR-TR concentration driven self-quenching is taking place.

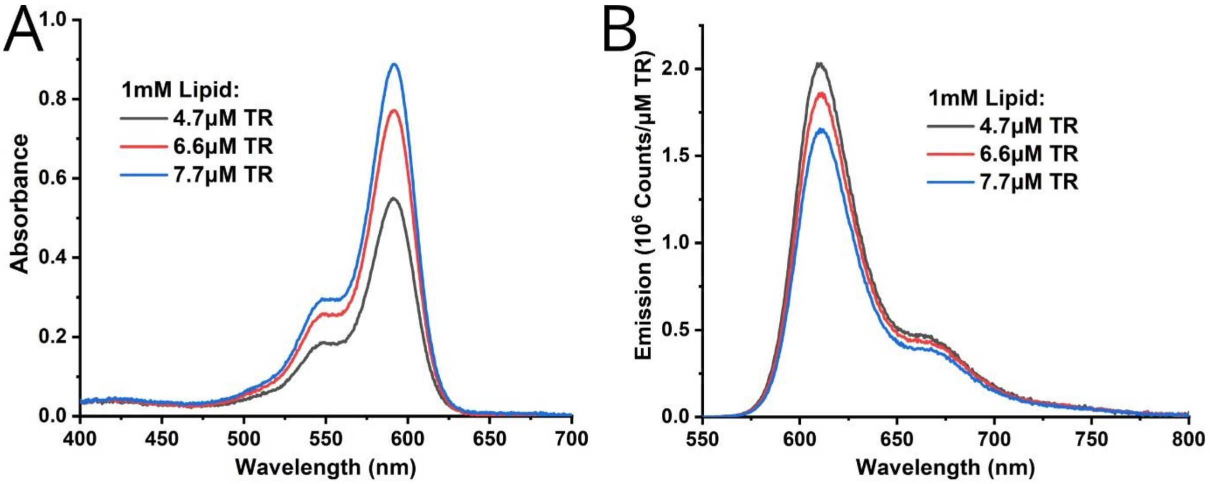


**Fig. S3** *Spectral evidence of concentration dependant TR fluorescence emission quenching.*

| Sample Name | TR Abs. | TR Conc. µM | TR Em. Counts | TR Em./ Abs Counts/ µM | Rel. TR Em. % |
|---|---|---|---|---|---|
| 5µM TR* | 0.550 | 4.7 | 9.62E+06 | 2.03E+06 | 100 |
| 7µM TR* | 0.771 | 6.6 | 1.23E+07 | 1.85E+06 | 91 |
| 8µM TR* | 0.888 | 7.7 | 1.25E+07 | 1.63E+06 | 80 |

**Table. S2** *Spectral data for demonstrating concentration dependant TR quenching*
*TR Abs./ Em. = TR fluorescence emission peak maxima divided by TR concentration in the sample.*
*Rel. Em. = TR fluorescence emission relative to Abs./Em. In the lowest concentration TR sample.*
*Samples were diluted before cuvette-based spectroscopy by the dilution factors indicated by * = x15.*

**ESI 5 - Analysis of emission spectra and calculation of relative emission enhancement for each dye-protein combination**

The relative fluorescence emission intensities of LH protein complexes were determined by measuring the protein's emission peak maximum in samples with varying concentrations of dye, under selective dye excitation (as described in main text **Methods** section 2.5). The values of LH protein fluorescence emission were corrected by subtracting the known overlap of the dye emission spectrum into the LH protein peak. At its emission peak at ~681nm LHCII has a contribution of 17.3 % and 6.5 % from TR and DiI relative to their emission maxima at ~610 nm and ~575 nm, respectively. At its emission peak at ~860 nm LH2 has a contribution of 0.5%, 13.7 %, and 6.7% from ATTO, Cy7, and DiR from their emission maxima at ~665 nm, ~790 nm, and ~785 nm, respectively. The ratio of emission/absorbance intensity (Em./Abs.) was calculated, in order to take into account slight variations in protein concentration between samples, and to allow a fair comparison of protein emission for a single sample set.

Em./Abs. was calculated by dividing the measured emission peak maxima by the µM concentration of protein in each sample (as calculated in **Table S1**).

The "Relative Emission" for LH protein complexes was calculated as:

$$Rel.Em. \ = \left(\frac{Em./Abs_{Dye}}{Em./Abs_{No\ dye}}\right) \times 100$$

from the Em./Abs. values of each sample with ($Em./Abs_{Dye}$) and without ($Em./Abs_{No\ Dye}$) dye. A value of 100% represents the level of fluorescence emission for a sample without any additional dye allowing us to assess the dye's effectiveness.

| Sample<br>Name (initial conc.) | TR Em.<br>Counts | LHCII Em.<br>Counts | Corr. LHCII Em.<br>Counts | LHCII Em./ Abs<br>Counts/ µM | LHCII Rel. Em.<br>% | Dye:LHCII Ratio | Dye:Chl Ratio |
|---|---|---|---|---|---|---|---|
| 0.7µM LHCII* | 3.4E+03 | 2.0E+06 | 2.0E+06 | 2.9E+06 | 100 | 0 | 0 |
| 0.7µM LHCII + 15µM TR* | 4.8E+05 | 3.8E+06 | 3.7E+06 | 6.0E+06 | 205 | 13.7 | 0.33 |
| 0.7µM LHCII + 22.5µM TR** | 6.9E+05 | 4.5E+06 | 4.3E+06 | 7.6E+06 | 259 | 24.6 | 0.59 |
| 0.7µM LHCII + 30µM TR** | 7.6E+05 | 5.2E+06 | 5.0E+06 | 8.3E+06 | 283 | 30.5 | 0.73 |
| 0.7µM LHCII + 37.5µM TR** | 8.2E+05 | 5.5E+06 | 5.4E+06 | 8.8E+06 | 302 | 36.8 | 0.88 |
| 0.7µM LHCII + 75µM TR*** | 1.1E+06 | 6.4E+06 | 6.3E+06 | 1.1E+07 | 361 | 76.5 | 1.82 |

| Sample<br>Name (initial conc.) | DiI Em.<br>Counts | LHCII Em.<br>Counts | Corr. LHCII Em.<br>Counts | LHCII Em./ Abs<br>Counts/ µM | LHCII Rel. Em.<br>% | Dye:LHCII Ratio | Dye:Chl Ratio |
|---|---|---|---|---|---|---|---|
| 0.7µM LHCII* | 1.3E+04 | 2.5E+05 | 2.5E+05 | 6.2E+05 | 100 | 0 | 0 |
| 0.7µM LHCII + 8µM* | 6.1E+04 | 3.3E+05 | 3.2E+05 | 8.7E+05 | 139 | 14.8 | 0.35 |
| 0.7µM LHCII + 12µM DiI* | 8.5E+04 | 3.8E+05 | 3.8E+05 | 1.1E+06 | 169 | 21.8 | 0.52 |
| 0.7µM LHCII + 20µM DiI* | 1.3E+05 | 4.4E+05 | 4.3E+05 | 1.3E+06 | 214 | 38.4 | 0.91 |
| 0.7µM LHCII + 10µM DiI* | 4.5E+04 | 3.3E+05 | 3.3E+05 | 1.0E+06 | 162 | 13.5 | 0.32 |
| 0.7µM LHCII + 20µM DiI** | 9.0E+04 | 4.0E+05 | 3.9E+05 | 1.2E+06 | 191 | 30.7 | 0.73 |
| 0.7µM LHCII + 30µM DiI** | 1.3E+05 | 4.9E+05 | 4.8E+05 | 1.6E+06 | 252 | 45.2 | 1.08 |
| 0.7µM LHCII + 40µM DiI** | 1.7E+05 | 5.1E+05 | 5.0E+05 | 1.7E+06 | 270 | 65.4 | 1.56 |
| 0.7µM LHCII + 50µM DiI** | 2.4E+05 | 5.5E+05 | 5.3E+05 | 1.9E+06 | 310 | 90.6 | 2.16 |
| 0.7µM LHCII + 75µM DiI*** | 2.3E+05 | 8.1E+05 | 7.9E+05 | 2.4E+06 | 384 | 135.2 | 3.22 |
| 0.7µM LHCII + 100µM DiI*** | 2.6E+05 | 9.1E+05 | 8.9E+05 | 2.8E+06 | 450 | 183.8 | 4.38 |

| Sample<br>Name (initial conc.) | ATTO Em.<br>Counts | LH2 Em.<br>Counts | Corr. LH2 Em.<br>Counts | LH2 Em./ Abs<br>Counts/ µM | LH2 Rel. Em.<br>% | Dye:LH2 Ratio | Dye:BChl Ratio |
|---|---|---|---|---|---|---|---|
| 0.5µM LH2* | 9.80E+02 | 1.27E+06 | 1.27E+06 | 3.02E+06 | 100 | 0 | 0 |
| 0.5µM LH2 + 2µM ATTO* | 1.77E+07 | 2.35E+06 | 2.26E+06 | 4.28E+06 | 142 | 2.6 | 0.1 |
| 0.5µM LH2 + 4µM ATTO* | 4.11E+07 | 2.69E+06 | 2.47E+06 | 4.95E+06 | 164 | 6.2 | 0.23 |
| 0.5µM LH2 + 6µM ATTO* | 7.96E+07 | 4.65E+06 | 4.23E+06 | 9.32E+06 | 309 | 13.2 | 0.49 |
| 0.5µM LH2 + 8µM ATTO* | 6.38E+07 | 4.37E+06 | 4.03E+06 | 8.36E+06 | 277 | 10.2 | 0.38 |
| 0.5µM LH2 + 10µM ATTO* | 1.04E+08 | 6.08E+06 | 5.52E+06 | 1.18E+07 | 391 | 17.8 | 0.66 |
| 0.5µM LH2 + 12µM ATTO* | 1.07E+08 | 6.27E+06 | 5.69E+06 | 1.23E+07 | 409 | 18.8 | 0.7 |

| Sample<br>Name (initial conc.) | Cy7 Em.<br>Counts | LH2 Em.<br>Counts | Corr. LH2 Em.<br>Counts | LH2 Em./ Abs<br>Counts/ µM | LH2 Rel. Em.<br>% | Dye:LH2 Ratio | Dye:BChl Ratio |
|---|---|---|---|---|---|---|---|
| 0.5µM LH2* | 4.22E+05 | 1.35E+07 | 1.34E+07 | 2.33E+07 | 100 | 0 | 0 |
| 0.5µM LH2 + 1.5µM Cy7* | 1.12E+07 | 1.74E+07 | 1.58E+07 | 3.24E+07 | 139 | 2.2 | 0.08 |
| 0.5µM LH2 + 3.0µM Cy7* | 2.65E+07 | 2.42E+07 | 2.05E+07 | 4.66E+07 | 200 | 4.6 | 0.17 |
| 0.5µM LH2 + 4.5µM Cy7* | 3.60E+07 | 3.30E+07 | 2.81E+07 | 6.10E+07 | 262 | 6.2 | 0.23 |
| 0.5µM LH2 + 6.0µM Cy7* | 5.34E+07 | 4.68E+07 | 3.95E+07 | 7.09E+07 | 304 | 8.4 | 0.31 |
| 0.5µM LH2 + 7.5µM Cy7* | 6.49E+07 | 5.26E+07 | 4.37E+07 | 7.93E+07 | 341 | 9.9 | 0.37 |
| 0.5µM LH2 + 9.0µM Cy7* | 7.30E+07 | 5.73E+07 | 4.74E+07 | 9.80E+07 | 421 | 12.9 | 0.48 |

| Sample<br>Name (initial conc.) | DiR Em.<br>Counts | LH2 Em.<br>Counts | Corr. LH2 Em.<br>Counts | LH2 Em./ Abs<br>Counts/ µM | LH2 Rel. Em.<br>% | Dye:LH2 Ratio | Dye:BChl Ratio |
|---|---|---|---|---|---|---|---|
| 0.5µM LH2 + 0µM DiR* | 4.58E+05 | 1.00E+07 | 9.97E+06 | 2.35E+07 | 100 | 0 | 0 |
| 0.5µM LH2 + 4µM DiR* | 1.13E+06 | 1.15E+07 | 1.14E+07 | 2.60E+07 | 110 | 1.3 | 0.05 |
| 0.5µM LH2 + 8µM DiR* | 1.54E+06 | 1.20E+07 | 1.19E+07 | 2.70E+07 | 115 | 2 | 0.07 |
| 0.5µM LH2 + 12µM DiR* | 1.90E+06 | 1.26E+07 | 1.25E+07 | 2.84E+07 | 120 | 2.7 | 0.1 |
| 0.5µM LH2 + 16µM DiR* | 2.64E+06 | 1.39E+07 | 1.37E+07 | 3.12E+07 | 133 | 3.9 | 0.15 |
| 0.5µM LH2 + 20µM DiR* | 3.13E+06 | 1.52E+07 | 1.50E+07 | 3.44E+07 | 146 | 4.8 | 0.18 |
| 0.5µM LH2 + 30µM DiR* | 1.08E+06 | 1.22E+07 | 1.21E+07 | 2.37E+07 | 101 | 6.3 | 0.23 |
| 0.5µM LH2 + 40µM DiR* | 1.26E+06 | 1.26E+07 | 1.25E+07 | 2.41E+07 | 102 | 7.8 | 0.29 |
| 0.5µM LH2 + 50µM DiR* | 2.53E+06 | 1.53E+07 | 1.52E+07 | 2.91E+07 | 124 | 10.3 | 0.38 |
| 0.5µM LH2 + 60µM DiR* | 3.45E+06 | 1.69E+07 | 1.67E+07 | 3.18E+07 | 135 | 12.4 | 0.46 |
| 0.5µM LH2 + 70µM DiR* | 4.25E+06 | 1.81E+07 | 1.78E+07 | 3.39E+07 | 144 | 14.6 | 0.54 |
| 0.5µM LH2 + 80µM DiR* | 4.37E+06 | 1.87E+07 | 1.84E+07 | 3.46E+07 | 147 | 16 | 0.59 |
| 0.5µM LH2 + 90µM DiR* | 4.73E+06 | 1.91E+07 | 1.88E+07 | 3.49E+07 | 148 | 18.1 | 0.67 |
| 0.5µM LH2 + 100µM DiR* | 5.40E+06 | 2.04E+07 | 2.01E+07 | 3.72E+07 | 158 | 20.4 | 0.75 |

**Table S3** *Emission spectra data and calculated relative enhancement of LH proteins.*

*Em. = Emission peak maxima for dye or LH protein.*

*Corr. Em. = Fluorescence emission peak maxima for dye or LH protein corrected for spectral overlap.*

*Abs./ Em. = Corrected LH protein fluorescence emission peak maxima divided by the protein concentration in the sample.*

*Rel. Em. = Relative LH protein emission (as defined by the equation above).*

*Samples were diluted before cuvette-based spectroscopy by the dilution factors indicated:*
** = x15, ** = x30, *** = x60*

**ESI 6 – Energy Transfer Efficiency (ETE) calculations from "1–T" vs. excitation spectra**

Before quantification of dye-to-protein ETE the dye peaks within the linear absorption and excitation spectra were decomposed from the measured (dye+protein) spectra to allow an accurate calculated area attributed to the signal from the dye alone. To do this, a representative LHCII or LH2 only spectrum of proteoliposomes (without dyes) was normalised to the peaks for the LH protein absorption in the combined sample and subtracted. This produced the dye-only spectra as shown in **Fig. S4.**

Comparison between fluorescence excitation spectra and linear absorption ("1–Transmission") spectra can indicate the connectivity of chromophores within an LH protein. We would expect perfectly overlapping excitation and 1–T spectra if all chromophores were well-connected as all energy absorbed by the system would manifest as fluorescence from the lowest energy (B)Chl emitter pigments. Comparison of fluorescence excitation and absorption spectra can also be used to determine the energy transfer efficiency between donor and acceptor molecules which undergo FRET. These two types of spectra were collected for LH proteoliposome samples with and without additional dyes. Linear absorption spectra were calculated as one minus the sample transmission (1 – T). Sample transmission, T, is defined as the fraction of the incident beam intensity which is transmitted through the sample $(I_T/I_0)$ and can be obtained from a sample's absorbance spectrum using the relationship:

$$T = \frac{I_T}{I_0} = 10^{-Abs}$$

where the logarithmic absorbance, Abs, is the standard measurement made by the instrument:

$$Abs = log_{10}\left(\frac{I_T}{I_0}\right)$$

If the excitation spectrum (*solid lines* in **Fig. S4**) shows lower intensity than the linear absorption (*dashed lines* in **Fig. S4**) there is less than 100% energy transfer between excitation donors and acceptors at that wavelength. The ratio of fluorescence excitation to linear absorption over the donor wavelength range is therefore directly related to ETE, which can be calculated as follows:

$$ETE = 1 - \frac{(1 - T_D) - Ex_D}{(1 - T_D)}$$

where $T_D$ = Donor transmission maximum; $Ex_D$ = Donor excitation maximum.

ETE calculations for selected samples are displayed in **Table S4**.

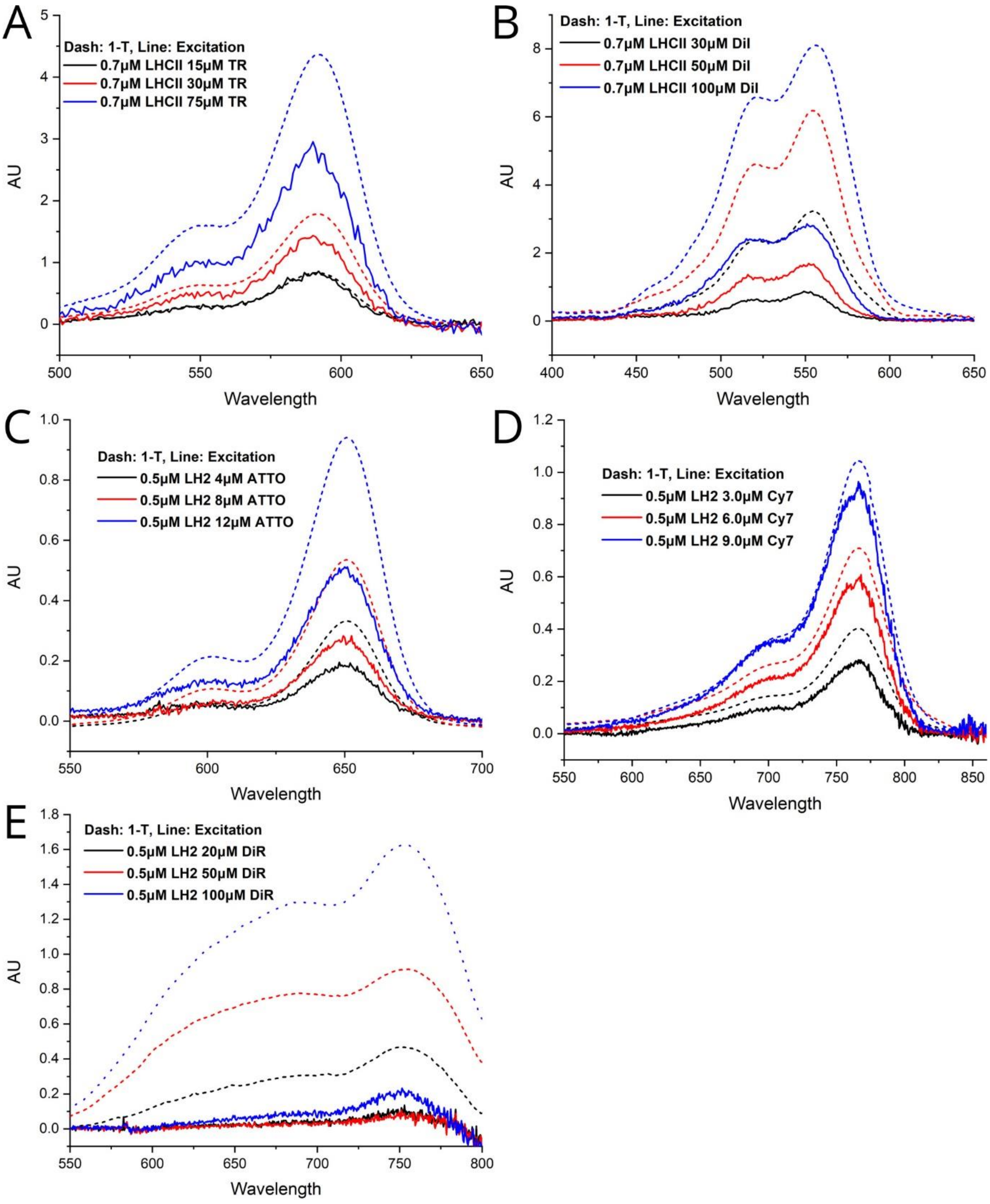


**Fig. S4.** *Linear absorption and fluorescence excitation spectra for ETE calculations on selected samples (LH component removed).*

| Sample<br>Name (initial conc.) | Range<br>(nm) | 1–T dye area | Ex dye area | ETE<br>(%) |
|---|---|---|---|---|
| 0.7µM LHCII + 15µM TR | 500-650 | 42 | 40 | 97 |
| 0.7µM LHCII + 30µM TR | 500-650 | 90 | 68 | 76 |
| 0.7µM LHCII + 75µM TR | 500-650 | 225 | 136 | 60 |
| 0.7µM LHCII + 30µM DiI | 450-650 | 229 | 57 | 25 |
| 0.7µM LHCII + 50µM DiI | 450-650 | 458 | 113 | 25 |
| 0.7µM LHCII + 100µM DiI | 450-650 | 670 | 218 | 33 |
| 0.5µM LH2 + 4µM ATTO | 580-680 | 12 | 8 | 66 |
| 0.5µM LH2 + 8µM ATTO | 580-680 | 21 | 11 | 53 |
| 0.5µM LH2 + 12µM ATTO | 580-680 | 39 | 22 | 57 |
| 0.5µM LH2 + 3.0µM Cy7 | 700-800 | 24 | 16 | 66 |
| 0.5µM LH2 + 6.0µM Cy7 | 700-800 | 44 | 36 | 81 |
| 0.5µM LH2 + 9.0µM Cy7 | 700-800 | 63 | 58 | 91 |
| 0.5µM LH2 + 20µM DiR | 550-850 | 63 | 7 | 11 |
| 0.5µM LH2 + 50µM DiR | 550-850 | 166 | 5 | 3 |
| 0.5µM LH2 + 100µM DiR | 550-850 | 275 | 15 | 5 |

**Table S4.** *Linear absorption spectra area, fluorescence excitation spectra area, and calculated ETE for selected samples. Abbreviations:*

*1-T = integrated area measured under linear absorption spectrum over the specified Range;*

*Ex. = integrated area measured under the excitation spectrum over the specified Range;*

*ETE = Energy transfer efficiency, as defined in the equation above.*

#### ESI 7 - Excitation spectra for LHCII proteoliposomes

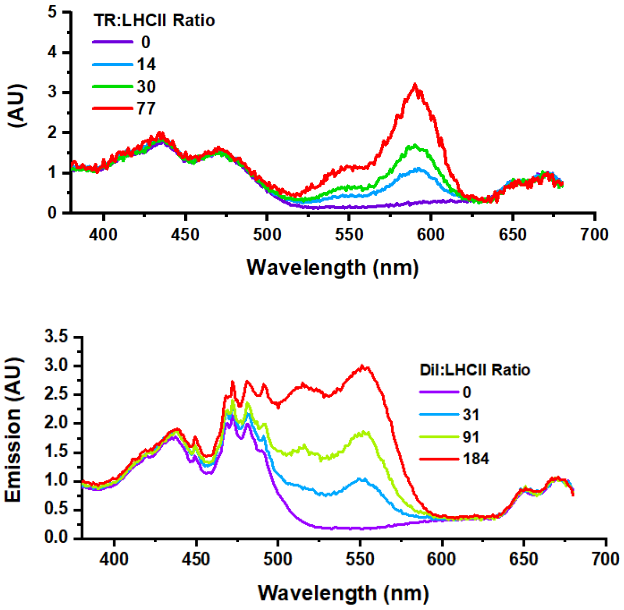


**Fig. S5.** *Comparison of the relative enhancement of plant LHCII by assessment of fluorescence excitation spectra. All spectra were acquired with the selective detection of LHCII emission at 686 nm, em/ex bandwidths 1/2 nm. Fluorescence excitation spectra for LHCII proteoliposomes containing either (A) TR-lipids, (B) Injected DiI.*

#### ESI 8 – Calculation of the "effective absorption strength" from excitation spectra

| Sample<br>Name (initial conc.) | Dye:<br>(B)Chl<br>ratio | Range 1 | Area 1 | Rel.<br>En. 1<br>(%) | Range 2 | Area 2 | Rel.<br>En. 2<br>(%) | Corr.<br>Rel. En.<br>(%) |
|---|---|---|---|---|---|---|---|---|
| 0.7μM LHCII | **0.00** | 380-680 | 225 | 100 | 500-650 | 41 | 100 | **100** |
| 0.7μM LHCII + 15μM TR | **0.33** | 380-680 | 273 | 122 | 500-650 | 81 | 198 | **118** |
| 0.7μM LHCII + 22.5μM TR | **0.59** | 380-680 | 282 | 126 | 500-650 | 98 | 239 | **125** |
| 0.7μM LHCII + 30μM TR | **0.73** | 380-680 | 292 | 130 | 500-650 | 107 | 260 | **129** |
| 0.7μM LHCII + 37.5μM TR | **0.88** | 380-680 | 322 | 143 | 500-650 | 124 | 303 | **137** |
| 0.7μM LHCII + 75μM TR | **1.82** | 380-680 | 370 | 165 | 500-650 | 177 | 431 | **160** |
| 0.7μM LHCII | **0.00** | 380-680 | 238 | 100 | 500-650 | 47 | 100 | **100** |
| 0.7μM LHCII + 8μM | **0.35** | 380-680 | 251 | 105 | 500-650 | 60 | 128 | **105** |
| 0.7μM LHCII + 12μM DiI | **0.52** | 380-680 | 267 | 112 | 500-650 | 70 | 149 | **110** |
| 0.7μM LHCII + 20μM DiI | **0.91** | 380-680 | 290 | 122 | 500-650 | 90 | 190 | **118** |
| 0.7μM LHCII + 10μM DiI | **0.32** | 380-680 | 247 | 104 | 500-650 | 57 | 120 | **104** |
| 0.7μM LHCII + 20μM DiI | **0.73** | 380-680 | 278 | 117 | 500-650 | 76 | 160 | **112** |
| 0.7μM LHCII + 30μM DiI | **1.08** | 380-680 | 301 | 126 | 500-650 | 95 | 202 | **120** |
| 0.7μM LHCII + 40μM DiI | **1.56** | 380-680 | 315 | 132 | 500-650 | 113 | 239 | **128** |
| 0.7μM LHCII + 50μM DiI | **2.16** | 380-680 | 356 | 149 | 500-650 | 144 | 305 | **141** |
| 0.7μM LHCII + 75μM DiI | **3.22** | 380-680 | 373 | 157 | 500-650 | 165 | 350 | **150** |
| 0.7μM LHCII + 100μM DiI | **4.38** | 380-680 | 466 | 196 | 500-650 | 231 | 489 | **177** |
| 0.5μM LH2 | **0.00** | 450-875 | 118 | 100 | 600-800 | 28 | 100 | **100** |
| 0.5μM LH2 + 2μM ATTO | **0.10** | 450-875 | 127 | 108 | 600-800 | 32 | 115 | **104** |
| 0.5μM LH2 + 4μM ATTO | **0.23** | 450-875 | 130 | 111 | 600-800 | 36 | 128 | **107** |
| 0.5μM LH2 + 6μM ATTO | **0.49** | 450-875 | 134 | 114 | 600-800 | 42 | 148 | **112** |
| 0.5μM LH2 + 8μM ATTO | **0.38** | 450-875 | 126 | 107 | 600-800 | 37 | 132 | **108** |
| 0.5μM LH2 + 10μM ATTO | **0.66** | 450-875 | 142 | 121 | 600-800 | 47 | 168 | **116** |
| 0.5μM LH2 + 12μM ATTO | **0.70** | 450-875 | 143 | 122 | 600-800 | 48 | 171 | **117** |
| 0.5μM LH2 | **0.00** | 450-875 | 115 | 100 | 600-800 | 27 | 100 | **100** |
| 0.5μM LH2 + 1.5μM Cy7 | **0.08** | 450-875 | 126 | 109 | 600-800 | 37 | 139 | **109** |
| 0.5μM LH2 + 3.0μM Cy7 | **0.17** | 450-875 | 134 | 117 | 600-800 | 47 | 177 | **118** |
| 0.5μM LH2 + 4.5μM Cy7 | **0.23** | 450-875 | 146 | 127 | 600-800 | 58 | 219 | **128** |
| 0.5μM LH2 + 6.0μM Cy7 | **0.31** | 450-875 | 163 | 141 | 600-800 | 72 | 270 | **139** |
| 0.5μM LH2 + 7.5μM Cy7 | **0.37** | 450-875 | 170 | 148 | 600-800 | 81 | 302 | **147** |
| 0.5μM LH2 + 9.0μM Cy7 | **0.48** | 450-875 | 194 | 169 | 600-800 | 101 | 378 | **164** |
| 0.5μM LH2 | **0.00** | 450-875 | 121 | 100 | 600-800 | 29 | 100 | **100** |
| 0.5μM LH2 + 6μM ATTO + 4.5μM Cy7 | **0.41** | 450-875 | 176 | 145 | 600-800 | 80 | 276 | **142** |
| 0.5μM LH2 + 6μM ATTO + 9μM Cy7 | **0.60** | 450-875 | 202 | 166 | 600-800 | 109 | 376 | **166** |
| 0.5μM LH2 + 12μM ATTO + 4.5μM Cy7 | **0.78** | 450-875 | 199 | 164 | 600-800 | 98 | 336 | **156** |
| 0.5μM LH2 + 12μM ATTO + 9μM Cy7 | **0.97** | 450-875 | 234 | 193 | 600-800 | 135 | 465 | **187** |
| 0.5μM LH2 + 0μM DiR | **0.00** | 450-875 | 112 | 100 | 600-800 | 26 | 100 | **100** |
| 0.5μM LH2 + 4μM DiR | **0.05** | 450-875 | 119 | 106 | 600-800 | 29 | 112 | **103** |
| 0.5μM LH2 + 8μM DiR | **0.07** | 450-875 | 110 | 99 | 600-800 | 28 | 107 | **102** |
| 0.5μM LH2 + 12μM DiR | **0.10** | 450-875 | 127 | 113 | 600-800 | 32 | 125 | **106** |
| 0.5μM LH2 + 16μM DiR | **0.15** | 450-875 | 117 | 105 | 600-800 | 31 | 122 | **105** |
| 0.5μM LH2 + 20μM DiR | **0.18** | 450-875 | 120 | 107 | 600-800 | 33 | 129 | **107** |
| 0.5μM LH2 + 30μM DiR | **0.23** | 450-875 | 115 | 103 | 600-800 | 28 | 108 | **102** |
| 0.5μM LH2 + 40μM DiR | **0.29** | 450-875 | 117 | 105 | 600-800 | 29 | 111 | **103** |
| 0.5μM LH2 + 50μM DiR | **0.38** | 450-875 | 117 | 105 | 600-800 | 31 | 121 | **105** |
| 0.5μM LH2 + 60μM DiR | **0.46** | 450-875 | 130 | 116 | 600-800 | 37 | 142 | **110** |
| 0.5μM LH2 + 70μM DiR | **0.54** | 450-875 | 132 | 118 | 600-800 | 39 | 151 | **112** |
| 0.5μM LH2 + 80μM DiR | **0.59** | 450-875 | 129 | 116 | 600-800 | 39 | 151 | **112** |
| 0.5μM LH2 + 90μM DiR | **0.67** | 450-875 | 131 | 117 | 600-800 | 41 | 157 | **113** |
| 0.5μM LH2 + 100μM DiR | **0.75** | 450-875 | 128 | 115 | 600-800 | 42 | 162 | **114** |

**Table S5** *Tabulated values for the calculation of "effective absorption enhancement" for all samples by integration of the area under excitation spectra. Notes:*

*Dye: (B)Chl ratio = repeated here, calculated in* ***Table S1****.*

*Range 1 = the spectral range chosen as the best representation of the entire proteins range (the full range that we collected data over here). Only the UV part of the protein's spectrum is excluded here due to instrument limitations.*

*Range 2 = the spectral range chosen as a good representation of the "spectral gap" that we are attempting to fill with the dye. This narrower range is an approximation of the "green spectral gap" of plant LHCII and the "red spectral gap" of bacterial LH2.*

*Area 1, Area 2 = the area measured by integration (using Origin Pro graphing software) using fluorescence excitation from main text* ***Figure 4*** *(for plant LHCII) and ESI* ***Figure S6*** *(for bacterial LH2). Area 1 or 2 is measured by using Range 1 or 2, respectively. The excitation spectra were normalised to an Emission intensity value of 1.00 at the (B)Chl $Q_y$ peak at 681nm (for LHCII) or 850nm (for LH2), as shown, before using the integration function.*

*Rel. En. 1 = Relative enhancement considering the full spectral range measured, in %:*

$$\frac{Area\ 1_{(dye+protein)}}{Area\ 1_{(protein)}} \times 100$$

*Rel. En. 2 = Relative enhancement considering the narrower range where the dye fills the protein spectral gap, in %:*

$$\frac{Area\ 2_{(dye+protein)}}{Area\ 2_{(protein)}} \times 100$$

*Corr. Rel. En. = These are the values plotted in main text* ***Figure 5****. The corrected values for the dye enhancement relative to entire spectral area (these values are similar to Rel. En. 2, but produce results that have lower noise):*

$$\left(\frac{Area\ 2_{(dye+protein)} - Area\ 2_{(protein)}}{Area\ 1_{(protein)}} + 1\right) \times 100$$

**ESI 9 - Evidence of ATTO-to-Cy7 energy transfer and increased effectiveness of dyes when working in conjunction**

Fluorescence emission spectra with selective ATTO excitation (625 nm) of liposome samples containing ATTO-lipids either with or without Cy7-lipid (**Fig. S6**). The emission spectra are normalised to the ATTO concentration to display the relative emission intensity of the ATTO dye. When Cy7 is included at a 6:5 ratio (ATTO:Cy7) there is a 95%+ reduction in ATTO relative emission (**Table S6**). This represents significant quenching of the fluorescence from ATTO is indicative of ATTO→Cy7 excitation energy transfer.

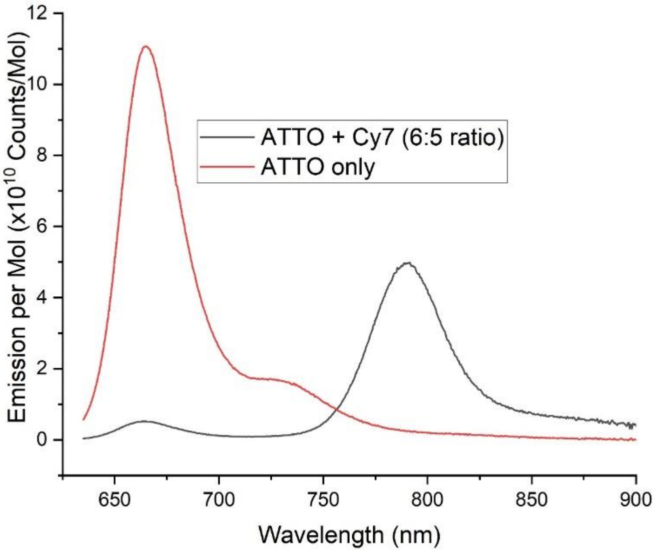


**Fig. S6** *Evidence of ATTO-to-Cy7 FRET from ATTO quenching. Fluorescence emission spectra of liposome samples with selective ATTO excitation (635 nm). Spectra are normalised to ATTO concentration calculated from absorption and molar extinction coefficient.*

| **Sample Name** | **ATTO Abs.** | **ATTO Abs Corr.** | **ATTO Conc. µM** | **Cy7 Abs.** | **Cy7 conc. µM** | **ATTO Em.** | **ATTO Em./Abs Counts/ µM** | **ATTO Rel. Em. %** | **ATTO:Cy7 Ratio** |
|---|---|---|---|---|---|---|---|---|---|
| 4µM ATTO* | 0.60 | 0.60 | 4.0 | 0.00 | 0.0 | 6.65E+07 | 1.65E+07 | 100.00 | 0.00 |
| 5µM ATTO + 6µM Cy7* | 1.17 | 0.91 | 6.0 | 1.25 | 5.0 | 4.71E+06 | 7.80E+05 | 4.70 | 1.21 |

**Table S6** *Spectral data for quantifying ATTO emission quenching due to ATTO-to-Cy7 energy transfer.*

*ATTO Abs. Corr. = ATTO absorption corrected for the spectral overlap of Cy7, ATTO has a contribution of 21% from Cy7 at its absorption maxima at ~647nm, relative to the Cy7 absorption maxima at ~750nm.*

*Abs./ Em. = ATTO fluorescence emission peak maxima divided by concentration in the sample.*

*Rel. Em. = ATTO fluorescence emission relative to Abs./Em. In the ATTO only sample.*

*Samples were diluted before cuvette-based spectroscopy by the dilution factors indicated:*
** = x15*

**ESI 10 – Numerical analysis of FLIM data investigating surface-supported membranes**

Crossover (spectral spillover) of signal between channels was calculated by measuring the image intensity of the dye-only (TR or DiI) or thylakoid-only control samples in all channels. These channel crossovers were used to correct intensity values for each combination of dye and hybrid thylakoid membrane.

| Sample | TR channel (mean counts) | Chl channel (mean counts) | Chl* channel (mean counts) | TR-to-Chl overlap (%) | TR-to-Chl* overlap (%) | Chl-to-Chl* overlap (%) |
|---|---|---|---|---|---|---|
| TR only | 44,130,191 | 30,949 | 1,344,423 | 0.07 | 3.05 | N/A |
| Chl only | 113,541 | 1,081,943 | 374,441 | N/A | 10.49 | 34.61 |

**Table S7** *Crossover controls for thylakoid membrane enhancement with TR.*

| Sample | DiI channel (mean counts) | Chl channel (mean counts) | Chl* channel (mean counts) | DiI-to-Chl overlap (%) | DiI-to-Chl* overlap (%) | Chl-to-Chl* overlap (%) |
|---|---|---|---|---|---|---|
| DiI only | 1,224,252 | 7,144 | 33,840 | 2.76 | 0.58 | N/A |
| Chl only | 8,082 | 318,361 | 113,364 | N/A | 2.54 | 35.60 |

**Table S8** *Crossover controls for thylakoid membrane enhancement with DiI.*

Tabulated data of the enhancement of hybrid membranes by TR or DiI:

| **Sample** | **Chl channel** | **Chl→Chl* crossover** | **TR channel** | **TR→Chl* crossover** | **Chl* channel** | **Chl* after crossover corrections** | **Relative Chl* intensity** |
|---|---|---|---|---|---|---|---|
| | $I_{Chl}$ | $I_{Chl\text{-}Chl*}$ | $I_{TR}$ | $I_{TR\text{-}Chl*}$ | $I_{Chl*}$ | $I_{Chl*-Corr.}$ | $Chl*_{Rel}$ |
| TR-DOPC 1 | 2.46E+04 | 8.52E+03 | 3.85E+07 | 1.17E+06 | 8.89E+05 | -2.84E+05 | N/A |
| TR-DOPC 2 | 1.99E+04 | 6.89E+03 | 3.13E+07 | 9.54E+05 | 8.08E+05 | -1.46E+05 | N/A |
| TR-DOPC 3 | 4.58E+04 | 1.58E+04 | 6.32E+07 | 1.93E+06 | 2.50E+06 | 5.73E+05 | N/A |
| TR-DOPC 4 | 3.35E+04 | 1.16E+04 | 4.35E+07 | 1.32E+06 | 1.18E+06 | -1.43E+05 | N/A |
| **Average** | **3.09E+04** | **1.07E+04** | **4.41E+07** | **1.34E+06** | **1.34E+06** | **0.00E+00** | **N/A** |
| *Error* | *5.69E+03* | *1.70E+03* | *6.83E+06* | *1.80E+05* | *3.93E+05* | *1.68E+05* | *N/A* |
| Hybrid (no TR) 1 | 1.10E+06 | 3.80E+05 | 1.19E+05 | 3.62E+03 | 3.18E+05 | -6.19E+04 | 84 |
| Hybrid (no TR) 2 | 1.31E+06 | 4.53E+05 | 1.47E+05 | 4.48E+03 | 5.21E+05 | 6.84E+04 | 115 |
| Hybrid (no TR) 3 | 9.81E+05 | 3.40E+05 | 9.65E+04 | 2.94E+03 | 2.79E+05 | -6.03E+04 | 82 |
| Hybrid (no TR) 4 | 9.39E+05 | 3.25E+05 | 9.18E+04 | 2.80E+03 | 3.79E+05 | 5.37E+04 | 117 |
| **Average** | **1.08E+06** | **3.74E+05** | **1.14E+05** | **3.46E+03** | **3.74E+05** | **0.00E+00** | **0** |
| *Error* | *8.29E+04* | *2.87E+04* | *1.26E+04* | *3.84E+02* | *5.31E+04* | *3.07E+04* | *8* |
| Hybrid with TR 1 | 3.53E+06 | 2.25E+05 | 6.51E+05 | 1.08E+05 | 5.35E+05 | 2.02E+05 | 190 |
| Hybrid with TR 2 | 3.78E+06 | 2.96E+05 | 8.56E+05 | 1.15E+05 | 6.92E+05 | 2.80E+05 | 195 |
| Hybrid with TR 3 | 4.54E+06 | 3.84E+05 | 1.11E+06 | 1.38E+05 | 9.52E+05 | 4.30E+05 | 212 |
| Hybrid with TR 4 | 5.65E+06 | 3.53E+05 | 1.02E+06 | 1.72E+05 | 9.71E+05 | 4.45E+05 | 226 |
| **Average** | **4.38E+06** | **3.15E+05** | **9.09E+05** | **1.33E+05** | **7.87E+05** | **3.39E+05** | **206** |
| *Error* | *4.77E+05* | *3.49E+04* | *1.01E+05* | *1.45E+04* | *1.06E+05* | *5.90E+04* | *8* |

| Sample | Chl channel | Chl→Chl* crossover | DiI channel | DiI→Chl* crossover | Chl* channel | Chl* after crossover corrections | Relative Chl* intensity (%) |
|---|---|---|---|---|---|---|---|
| | $I_{Chl}$ | $I_{Chl\text{-}Chl^*}$ | $I_{DiI}$ | $I_{DiI\text{-}Chl^*}$ | $I_{Chl^*}$ | $I_{Chl^*-Corr.}$ | $Chl^*_{Rel}$ |
| Hybrid (no DiI) 1 | 2.71E+05 | 9.63E+04 | 9.74E+03 | 2.47E+02 | 1.05E+05 | 8.27E+03 | 109 |
| Hybrid (no DiI) 2 | 2.12E+05 | 7.54E+04 | 6.33E+03 | 1.61E+02 | 6.99E+04 | -5.65E+03 | 93 |
| Hybrid (no DiI) 3 | 3.54E+05 | 1.26E+05 | 8.20E+03 | 2.08E+02 | 1.12E+05 | -1.40E+04 | 89 |
| Hybrid (no DiI) 4 | 4.37E+05 | 1.56E+05 | 8.06E+03 | 2.05E+02 | 1.66E+05 | 1.05E+04 | 107 |
| **Average** | **3.18E+05** | **1.13E+05** | **8.08E+03** | **2.05E+02** | **1.13E+05** | **-2.05E+02** | **99** |
| *Error* | *4.92E+04* | *1.75E+04* | *6.96E+02* | *1.77E+01* | *1.99E+04* | *1.01E+04* | *9* |
| Hybrid (5% DiI) 1 | 4.44E+05 | 1.58E+05 | 7.17E+04 | 1.98E+03 | 2.42E+05 | 8.19E+04 | 152 |
| Hybrid (5% DiI) 2 | 2.17E+05 | 7.74E+04 | 6.47E+04 | 1.79E+03 | 1.17E+05 | 3.77E+04 | 149 |
| Hybrid (5% DiI) 3 | 3.40E+05 | 1.21E+05 | 6.49E+04 | 1.79E+03 | 1.84E+05 | 6.09E+04 | 150 |
| Hybrid (5% DiI) 4 | 2.41E+05 | 8.60E+04 | 7.06E+04 | 1.95E+03 | 1.10E+05 | 2.19E+04 | 126 |
| **Average** | **3.11E+05** | **1.11E+05** | **6.80E+04** | **1.88E+03** | **1.63E+05** | **5.06E+04** | **144** |
| *Error* | *5.17E+04* | *1.84E+04* | *1.85E+03* | *5.12E+01* | *3.11E+04* | *1.32E+04* | *6* |
| Hybrid (10% DiI) 1 | 3.49E+05 | 1.24E+05 | 1.83E+05 | 5.05E+03 | 1.97E+05 | 6.74E+04 | 154 |
| Hybrid (10% DiI) 2 | 2.46E+05 | 8.77E+04 | 1.26E+05 | 3.47E+03 | 1.77E+05 | 8.60E+04 | 198 |
| Hybrid (10% DiI) 3 | 2.78E+05 | 9.90E+04 | 1.22E+05 | 3.37E+03 | 1.61E+05 | 5.86E+04 | 159 |
| Hybrid (10% DiI) 4 | 2.46E+05 | 8.76E+04 | 1.80E+05 | 4.97E+03 | 1.49E+05 | 5.63E+04 | 164 |
| **Average** | **2.80E+05** | **9.96E+04** | **1.53E+05** | **4.22E+03** | **1.71E+05** | **6.71E+04** | **169** |
| *Error* | *2.42E+04* | *8.62E+03* | *1.66E+04* | *4.60E+02* | *1.03E+04* | *6.74E+03* | *10* |

**Table S9** *Analysis for calculating hybrid thylakoid membrane emission enhancement. All values are total number of counts measured in each corral.*

$I_{Chl}$, $I_{TR}$, $I_{DiI}$, $I_{Chl^*}$: *All intensity values are the averages calculated from the accumulated fluorescence counts in n = 4 corrals of hybrid membranes with imaging conditions as described in* ***Methods 2.7****.*

$I_{TR\text{-}Chl^*}$, $I_{DiI\text{-}Chl^*}$, $I_{Chl\text{-}Chl^*}$: *Crossover values are the contribution from different other components into the Chl* channel and are calculated by multiplying the intensity of each channel by the overlap factors shown in* ***Tables S7+S8****.*

$I_{Chl^*-Corr.}$: *Corrected Chl* intensity is calculated by subtracting the crossover contributions (*$I_{TR\text{-}Chl^*}$, $I_{DiI\text{-}Chl^*}$, $I_{Chl\text{-}Chl^*}$*) from the raw Chl* intensity (*$I_{Chl^*}$*).*

$Chl^*_{Rel}$ **:** *Relative Chl* intensity is the calculated as the intensity of the Chl* channel taking the crossover intensity into account, so that 100% represents the baseline emission from the Chl-proteins and values above 100% represent enhancement due to excitation energy transfer from TR or DiI, as follows:*

$$Chl^*_{Rel} = \left(\frac{I_{Chl^*-Corr.} + I_{Chl-Chl^*}}{I_{Chl-Chl^*}}\right) \times 100$$

**Calculation: amount of dye introduced into hybrid membranes**

Considering the samples shown in main text **Figure 6** and analysed by FLIM:

**(A) TR-lipids:**
0.5% TR-lipids were included in the DOPC liposomes, i.e., 1:2000 TR:DOPC mole-to-mole. These lipid vesicles merge with the thylakoids membranes during the formation of the supporting membrane termed "hybrid membranes". The protocol used a 3:1 ratio of liposomes to thylakoid membranes, and the thylakoid membranes are comprised of 60-70% protein by weight. It is not possible to know the final concentration of proteins and TR in the sample but our previous publication provided an estimate that the protein represented 1-3 % of the final membrane area [7] and so the TR-to-lipid ratio may be similar to the starting ratio 1:2000 of TR:DOPC.

**(B) DiI:**
DiI was injected into the droplet above the pre-formed hybrid membrane and it seemed logical that only a fraction of the dye would actually insert into the membrane. Therefore to insert a comparabale amount of DiI dye as for TR dye it seemed reasonable to inject an order of magnitude excess as compared to TR (5% rather than 0.5%). This was based upon the estimated number of lipid present in corrals:

- We estimated, 62,500 total corrals based on a 10×10 mm patterned area of the glass coverslip ($1\times10^8$ $\mu m^2$) and 2-D array pattern with one corral in every 40×40 µm region (1,600 $\mu m^2$).
- We estimated $1.106 \times 10^{-15}$ mole lipid per corral based on 20×20 µm corral dimensions ($4\times10^8$ $nm^2$) and a known area per lipid of 0.6 $nm^2$ and Avagadro's number ($6.022\times10^{23}$ $mol^{-1}$). We are only consider the top monolayer of lipids in this calculation (otherwise the number of lipids is double).
- Therefore, this is a total of $6.91 \times 10^{-11}$ mol total lipid (69.1 pmol) (62,500 x $1.106 \times 10^{-15}$)
- To achieve 5% or 10% DiI w.r.t total lipids: we calculated that $3.45 \times 10^{-12}$ mol DiI should be injected [ (5/100) × ($6.91 \times 10^{-11}$) mol ].
- This equates to 3.227 ng DiI based on a molecular mass of 933.87.
- Using a stock of DiI dissolved into ethanol at a concentration of 10 µg/mL (10 ng/uL), a volume of 0.32 µL or 0.64 µL was required.
- This volume was injected directly into the approx. 200uL droplet of aqueous buffer that immersed the hybrid membranes supported on the glass coverslip. This would produce a final concentration of ~17 uM or ~34 uM of DiI in the sample droplet.

After injecting this volume of DiI in ethanol, there was a 15 min incubation period where DiI molecules were expected to associate with the membranes. Subsequently, the sample was washed and then imaged by FLIM. DiI presumably inserted into the lipid bilayer and interacted with the thylakoid proteins during this time. It is not possible to calculate the actual amount of DiI the incorporated into these membranes, but we can quantify the change in the sample's fluorescence.

**Reference List**

[1] P.G. Adams, C. Vasilev, C.N. Hunter, M.P. Johnson, Correlated fluorescence quenching and topographic mapping of Light-Harvesting Complex II within surface-assembled aggregates and lipid bilayers, Biochimica et biophysica acta, 1859 (2018) 1075-1085, 10.1016/j.bbabio.2018.06.011.

[2] R.J. Porra, W.A. Thompson, P.E. Kriedemann, Determination of accurate extinction coefficients and simultaneous equations for assaying chlorophylls a and b extracted with four different solvents: verification of the concentration of chlorophyll standards by atomic absorption spectroscopy, Biochimica et biophysica acta, 975 (1989) 384-394, 10.1016/S0005-2728(89)80347-0.

[3] T. Förster, Delocalized excitation and excitation transfer, Modern Quantum Chemistry Istanbul Lectures, 3 (1965) 93-137,

[4] L.M. Loura, Simple estimation of Forster Resonance Energy Transfer (FRET) orientation factor distribution in membranes, Int. J. Mol. Sci., 13 (2012) 15252-15270, 10.3390/ijms131115252.

[5] T. Sahin, M.A. Harris, P. Vairaprakash, D.M. Niedzwiedzki, V. Subramanian, A.P. Shreve, D.F. Bocian, D. Holten, J.S. Lindsey, Self-Assembled Light-Harvesting System from Chromophores in Lipid Vesicles, The journal of physical chemistry. B, 119 (2015) 10231-10243, 10.1021/acs.jpcb.5b04841.

[6] J.A. Titus, R. Haugland, S.O. Sharrow, D.M. Segal, Texas red, a hydrophilic, red-emitting flourophore for use with flourescein in dual parameter flow microfluorometric and fluorescence microscopic studies, Journal of Immunological Methods, 50 (1982) 193-204, 10.1016/0022-1759(82)90225-3.

[7] S.A. Meredith, T. Yoneda, A.M. Hancock, S.D. Connell, S.D. Evans, K. Morigaki, P.G. Adams, Model lipid membranes assembled from natural plant thylakoids into 2D microarray patterns as a platform to assess the organization and photophysics of light-harvesting proteins, Small, 17 (2021) e2006608, 10.1002/smll.202006608.
